# Widespread coupling of promoters and terminators of transcription in bacteria

**DOI:** 10.64898/2026.08.19.745784

**Authors:** Andrew G. Fletcher, David Forrest, Charles Cooper, Matthew P. Adams, Achillefs N. Kapanidis, David C. Grainger

## Abstract

Transcription underpins the expression of genetic information and is viewed as a series of independent initiation and termination events. In bacteria, promoters and terminators of transcription are therefore considered distinct regulatory elements. Here, we challenge this view. Genome-scale analyses reveal that promoters and terminators are extensively coupled. Thus, initiation and termination sites, for adjacent transcription units, frequently overlap. This conserved organisation arises because the DNA sequences, which direct termination, also contribute to promoter function. This couples neighbouring transcription units and generates regulatory interference between RNA polymerases. These findings define a universal mechanism for coordinating transcription across bacterial genomes.

## INTRODUCTION

Promoters are DNA sequences that bind RNA polymerase to stimulate transcription^1^. Different promoter regions have distinct roles and, in bacteria, the housekeeping -10 element is key. This motif, with the consensus 5′-TATAAT-3′, named according to location, contacts the RNA polymerase σ subunit to drive DNA unwinding^2^. Additional sequences bind and correctly position RNA polymerase at the promoter. For instance, 5′-TTGACA-3′ motifs at position -35, extended -10 elements with an upstream 5′-TGn-3′ sequence, or similarly positioned T-tracts, can all contribute^3–6^. Once a promoter has been bound, and the DNA duplex unwound, RNA polymerase transcribes the DNA immediately downstream of the transcription start site (TSS, +1). The enzyme does not dissociate from the promoter to do this, instead pulling downstream DNA into the active centre. This process, called “DNA scrunching”, generates mechanical stress and a short RNA^7,8^. The acquired potential energy can drive promoter escape or, more frequently, transcription is aborted and the RNA released^7,8^. Repeating until transcription elongation begins, the process is usually rate limiting^9^. As such, RNA polymerase is typically bound at the promoter much longer than at any position in the downstream transcription unit. Indeed, for 23% of promoters, the RNA polymerase is “poised” and cannot enter the elongation phase of transcription^9^.

Comparatively, transcription elongation is rapid, with ∼45 nucleotides being added to the RNA chain every second ^10^. Even so, RNA polymerase occasionally stalls^11^. This can be due to RNA secondary structure, DNA sequence, positive supercoiling, or DNA bound proteins^12^. For example, an elongation complex cannot readily dislodge promoter-bound RNA polymerase^13^. Whilst most pauses are temporary, the phenomenon also precedes termination^14^. Intrinsic terminators are pause sequences with an inverted repeat that encodes an RNA stem-loop^15^. The hairpin forms in the transcript exit channel whilst RNA polymerase is stalled. Resulting structural changes disrupt the RNA:DNA hybrid, close to the enzyme active centre, and the nascent RNA is released^16^. This is made easier by weak inter-nucleic acid base pairing. Hence, stem-loop encoding motifs are followed by a non-template strand T-tract^14^. The transcription termination site (TTS) is found directly after this element. Like promoter escape, termination is comparatively slow^17^. Indeed, RNA polymerase is trapped at some terminators until dislodged by an elongation complex^18,19^. Intrinsic terminators differ in efficiency, and RNA polymerase may “read through” some such sites^20^.

Whole genome sequences, computational tools, and associated experimental approaches, have driven efforts to map all promoters and terminators. For example, sequencing can identify transcript ends, DNA motifs can be found bioinformatically, and conserved features are evident across species^21–24^. In this work, we used such approaches to map transcription initiation and termination in diverse bacteria. We show that promoters and terminators, on the same DNA strand, frequently co-locate. Specifically, TSSs frequently occur ∼10 bp downstream of TTSs. We refer to such terminators as being promoter coupled. At these loci, DNA sequences serve a dual purpose and terminator motifs encompass key promoter elements. Moreover, the proximity of termination and initiation sites creates regulatory interference. Most notably, coupled promoters increase termination efficiency. We argue that this is a universal mechanism used to control gene expression.

## RESULTS

### Promoters and terminators are separated by preferred distances in Acinetobacter baumannii

Patterns of transcription have been mapped for many bacteria. However, interplay between initiation and termination has received little attention^25–27^. In a search for such interactions, we mapped RNA 5’ and 3’ ends, using cappable-seq and term-seq respectively, in the human pathogen *Acinetobacter baumannii*^23,24^. Our analysis revealed 12,529 TSSs, and 1,810 TTSs (Tables S1 and S2). For each termination site, we determined the distance to the nearest promoter on the same DNA strand (Figure 1a). All terminators are within a few thousand base pairs of the nearest same-strand promoter. This is unsurprising, the average *A. baummanii* gene is 903 bp in length and most transcription units contain several genes^28^. Even so, some TTS-TSS pairs should be much closer. This is particularly true for transcription units in the same orientation (i.e. tandemly organised). Here, termination and promotion, of upstream and downstream transcription, must involve the same intergenic DNA. The mean length of such intergenic regions, in *A. baumannii*, is 142 bp. With this in mind, we examined the elevated signal at the centre of the Figure 1a heatmap (see expansion). Surprisingly, this identified 200 TTS within 35 bp of a downstream TSS, with the most common intervening distance being just 10 bp (Table S1). Hence, many terminators closely overlap a promoter. We refer to these terminators as promoter-coupled, and those not overlapping a promoter as distinct (Figure 1a, schematic).

**Figure 1.**
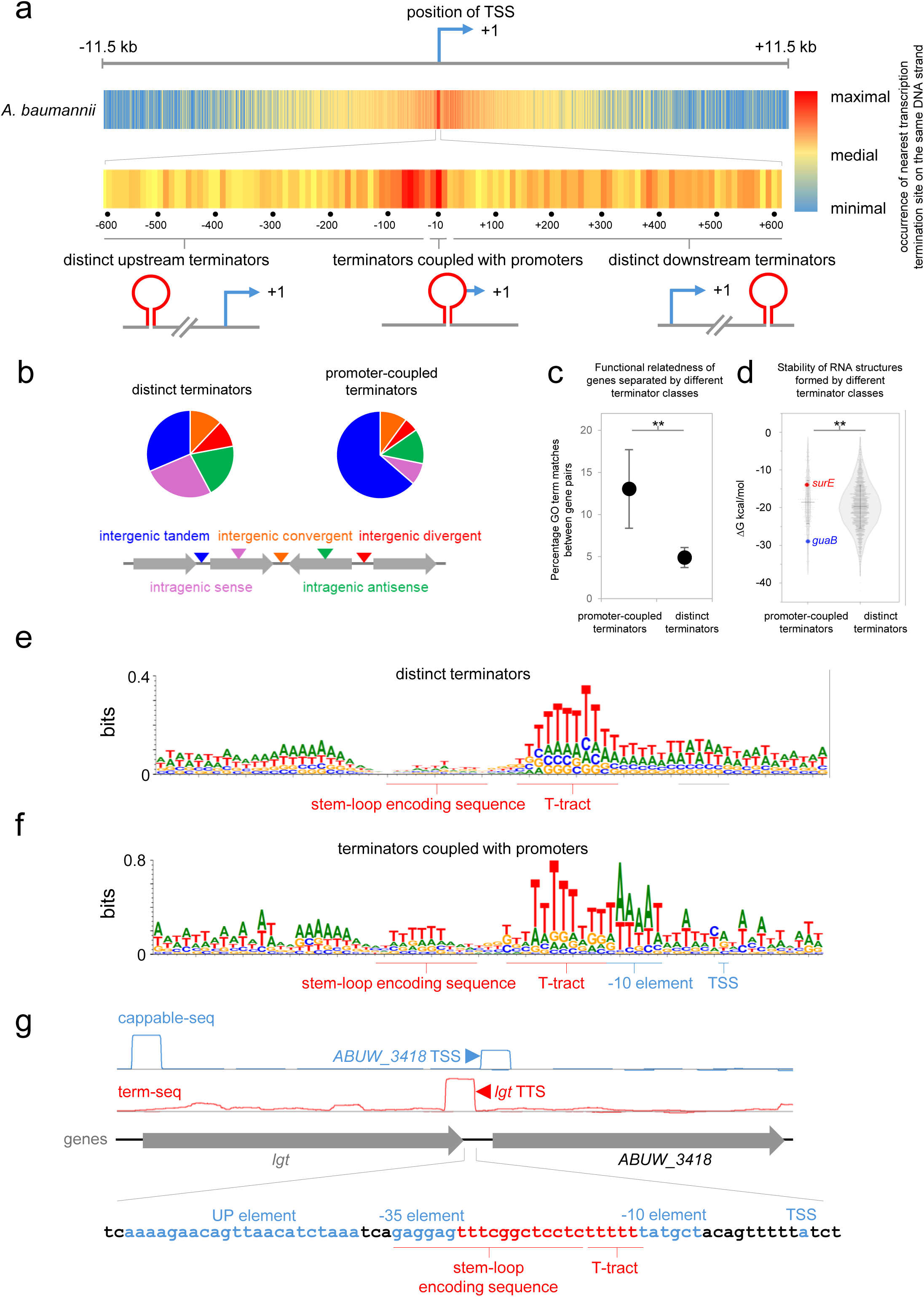
Widespread coupling of terminators and promoters in *Acinetobacter baumannii*. **a. Heatmaps showing the global distribution of TTSs with respect to TSSs in *A. baumannii*.** All TSSs are aligned at the centre of the heatmap (bent arrow, labelled +1) and colours indicate the abundance of TTSs, grouped in 100 bp bins, at the relevant position for the same DNA strand. The expansion shows 600 bp, either side of the aligned promoters, with the data grouped in 10 bp bins. In the schematic, terminators are shown as red lollipops and transcription start sites (labelled +1) as blue arrows. **b. Terminator location with respect to coding DNA sequences.** The pie charts show the percentage of distinct terminators (left) and those coupled to promoters (right) with respect to genes, in different orientations, as illustrated by the schematic. Genes are shown as arrows and triangles indicate the different location categories. **c. Functional relatedness of gene pairs flanking terminators.** Functional relatedness was determined by calculating the percentage of GO terms shared by gene pairs flanking an intergenic terminator. The data for all distinct or promoter-coupled terminators are presented as an interval plot. Error bars indicate the 95% confidence interval. P was determined using a two-tailed t-test, assuming unequal variance **d. Predicted RNA 3’-end folding stability.** Gibbs free energy changes associated with folding of the final 80 nucleotides of each RNA 3’-end identified by term-seq. The data for distinct and promoter-coupled terminators are separately grouped. Error bars show standard deviation, and the central horizontal line shows the mean. P was determined using a two-tailed t-test, assuming unequal variance **e,f. DNA sequence motifs associated with distinct terminators (e) or terminators coupled to promoters (f).** Terminator and promoter elements are underlined red and blue respectively. For panel e, we aligned sequences for all terminators not coupled to a promoter. The motif in panel f was generated by aligning all terminators where TTSs were between 8 and 11 bp upstream of a TSS (highlighted peak in Figure 5c). The grey underline shows a perfect match to a promoter -10 element at distinct terminators. This may be due to instances where a coupled promoter was inactive in our conditions. **g. An example of a promoter-coupled terminator in *A. baumannii*.** The TSS and TTS, identified by cappable-seq and term-seq, are indicated by blue and red triangles respectively. The expansion shows the associated DNA elements, also coloured blue or red. Key sections of the promoter and terminator are labelled.

### Promoter-coupled terminators are enriched between tandem genes with related functions

Locations of distinct and promoter-coupled terminators are summarised in Figure 1b. The latter are significantly more likely to be between tandem genes (P = 9.2 e^-9^, chi-squared test) and are seldom found in coding DNA. Further, the median intergenic region length, between tandem coding sequences, was 148 bp for distinct and 114 bp for promoter-coupled terminators. The difference approximates to the space reduction achieved if a promoter and terminator overlap. Shared Gene Ontology (GO) terms, for coding sequences flanking terminators, are in Figure 1c^29^. Functional similarity was higher for tandem gene pairs separated by promoter-coupled, rather than distinct, terminators. We conclude that the extrinsic context of distinct and promoter-coupled terminators is measurably different.

### Promoter-coupled terminators have distinct DNA sequence properties

We next turned our attention to inherent terminator properties. Transcript 3′-end structures were predicted using RNAFold, which also determines the Gibbs free energy change (ΔG) upon RNA folding^30^ (Figure 1d). Whilst ΔG distributions overlap, the average free energy change for folding of distinct terminators (-19.6 kcal/mol) is significantly larger than for promoter-coupled examples (-18.5 kcal/mol). Note that the range of ΔG predictions is large because stem-loop sequences are diverse^31^. Hence, upon generation of DNA sequence logos, the non-template strand T-tract is predominant (Figures 1e and 1f). Promoter-coupled terminators are distinguished by a -10 hexamer, downstream of the terminator T-tract, and a purine at the expected TSS location (compare Figures 1e and 1f). We also observed a pyrimidine immediately upstream of the TSS, as is common in *A. baumannii*^27^. Note that both terminator classes have an upstream A-tract, and this is because some terminators act bidirectionally (Table S2)^18^.

### Coupling of promoters and terminators can be predicted de novo

We reasoned that *de novo* discovery of promoter-coupled terminators should be possible bioinformatically. Hence, a position weight matrix was used to select promoters, and terminators were found with TransTermHP (Tables S3 and S4)^21^. Distances between predicted TSSs, and the nearest same-strand TTS, are summarised for *A. baumannii* in Figure S1a. The result is consistent with experimental mapping of RNA 5′ and 3′ ends. Thus, sites for transcription termination and initiation, separated by ∼10 bp, are strongly over-represented. In this preferred configuration, some DNA sequences must be used for both initiation and termination. For instance, in the Figure 1g example, the promoter -35 and RNA stem-loop motifs coincide. Further, the terminator T-tract is positioned, just upstream of the -10 hexamer, to enhance promoter activity^32^.

### Dual function DNA elements promoter-coupled terminators

To understand the role of dual-purpose DNA elements, we made a semi-synthetic DNA sequence. Our starting point was the *guaB* terminator (Table S2). We expected this motif to end transcription efficiently as the predicted ΔG for RNA folding is large (Figure 1d, blue datapoint). Briefly, and guided by the motif in Figure 1f, the sequence was modified to include a -10 element immediately downstream of the terminator T-tract. In this context, the stem-loop encoding DNA also acts as the -35 hexamer. The sequence is shown in Figure 2a, alongside mutated derivatives. Critically, there is no upstream promoter. Hence, RNA polymerase is not delivered to the promoter-coupled sequence for termination, avoiding potential interference. Each sequence was cloned upstream of the λ*oop* terminator to provide a template for *in vitro* transcription (Figure 2b). Consistent with elements being dual purpose, mutations in the terminator T-tract, and stem-loop encoding DNA, reduced coupled-promoter activity (Figure 2b). Similar results were obtained *in vivo* using *lacZ* fusions (compare Figures 2b and 2c).

**Figure 2.**
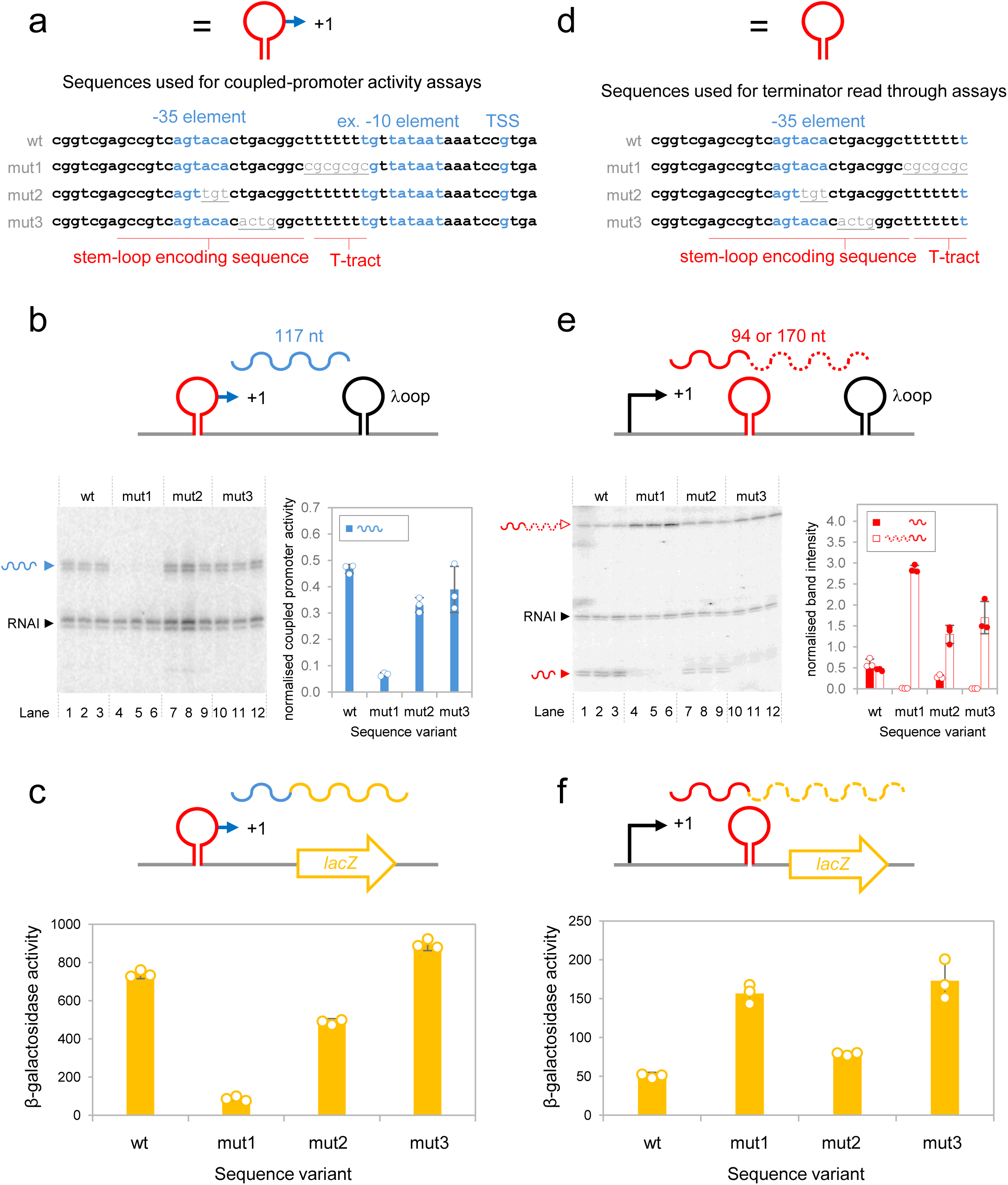
Dual purpose DNA elements permit coupling of promoters and terminators. **a. A semi-synthetic promoter-coupled terminator.** The wild type DNA sequence contains both the functional terminator (red underlining) and coupled promoter (blue text). Mutated derivatives have base changes shown in plain text and underlined. The schematic shows a graphic representation of the wild type sequence also used elsewhere in the figure, where the red lollipop indicates the DNA sequence encoding the stem-loop and the blue arrows shows the overlapping promoter. **b. Impact of mutations on coupled promoter activity *in vitro*.** The schematic illustrates the promoter-coupled terminator was cloned upstream of the λ*oop* terminator. The size of the expected RNA (wavy line) is indicated. There is no upstream promoter and hence no elongation complexes are expected to terminate at the cloned sequence. The gel image shows results of *in vitro* transcription assays using the DNA templates shown in panel a. Reactions were done in triplicate and the RNAI transcript, generated from the plasmid template replication origin, serves as an internal control. The bar chart shows mean production of the 117 nucleotide (nt) RNA from each template relative to the RNAI control. Errors bars show standard deviation (n=3) and the overlapping dot plot shows the data from each replicate. **c. Impact of mutations on coupled promoter activity *in vivo*.** The DNA fragments illustrated in panel a were cloned upstream of *lacZ* in plasmid pRW50 to create *lacZ* fusions (see schematic). Mean β-galactosidase activities of transformants are shown in the bar chart. Errors bars show standard deviation (n=3) and the overlapping dot plot shows the data from each replicate. **d. A truncated semi-synthetic promoter-coupled terminator.** The DNA sequences are the same as those shown in panel a except that they are truncated from the downstream end to remove the TSS and -10 element of the coupled promoter. The schematic shows the truncated wild type sequence, cloned upstream of the λ*oop* terminator. **e. Impact of mutations on coupled terminator activity *in vitro*.** The truncated sequences were cloned upstream between the λ*oop* terminator and a constitutive upstream promoter (see schematic, with expected size of terminated and read-through transcripts). The gel image shows results of an *in vitro* transcription assays using the DNA templates in panel d. Reactions were done in triplicate and the RNAI transcript, generated from the plasmid template replication origin, serves as an internal control. The bar chart shows mean production of the terminated (red bars) and read-through (open bars) RNA from each template, relative to the RNAI control. Errors bars show standard deviation (n=3) and the overlapping dot plot shows the data from each replicate. **f. Impact of mutations on terminator read through *in vivo*.** The DNA fragments illustrated in panel d were cloned upstream of *lacZ* in plasmid pRW50 to create *lacZ* fusions (see schematic). Mean β-galactosidase activities of transformants, arising from terminator read through, are shown in the bar chart. Errors bars show standard deviation (n=3) and the overlapping dot plot shows the data from each replicate.

To confirm that the same mutations also reduced termination, a shorter version of each fragment was used. Truncated sequences lack the coupled -10 motif to avoid promoter-bound RNA polymerase interference (Figure 2d). Expected transcripts are illustrated in Figure 2e, where terminated and read-through RNAs are 94 nt and 170 nt in length respectively. All mutations reduced terminator efficiency, *in vitro* and *in vivo*, as expected (Figures 2e and 2f).

### Coupled promoter activity modulates terminator efficiency in vitro

Beyond optimising use of intergenic space, promoter-terminator coupling could be regulatory. Specifically, because of the long dwell time, during initiation and termination, RNA polymerase interference could be key. We turned our attention to the *surE* terminator and coupled *nlpD* promoter (Figure 3a). This DNA region was cloned between the upstream *surE* promoter and λ*oop* terminator (Figure 3b schematic). The full base sequence is in Figure S2. All expected transcripts were detected following *in vitro* transcription (Figure 3b, lanes 1-3). However, the coupled *nlpD* promoter was poorly active and termination, at the overlapping *surE* terminator, was inefficient. The latter is consistent with unstable mRNA 3’-end folding (Figure 1d, red datapoint) but contrasts with the robust term-seq signal (Figure 3a). To understand any impact of RNA polymerase interference, at the promoter-coupled terminator, we first mutated the upstream *surE* promoter (Figure S2 and further Figure 3b schematics). As expected, the 94 nt and 207 nt transcripts were lost (lanes 4-6 and Figure 3c). Further, there was no change in *nlpD* transcription (lanes 4-6 and Figure 3d). Inactivation of the *nlpD* promoter increased read through of the coupled *surE* terminator by 58% (lanes 7-9 and Figure 3c). Conversely, when the *nlpD* promoter was improved, *surE* terminator read through decreased by 25% compared to the starting DNA (lanes 10-12 and Figure 3c). Hence, *nlpD* promoter activity modulates coupled *surE* terminator efficiency. In similar experiments, we also tested the semi-synthetic promoter-coupled terminator (Figure S3). Here, termination was very efficient, and the coupled promoter much more active (Figure S3b, lanes 1-3). Strikingly, mutation of the coupled promoter substantially increased terminator readthrough (Figure S3b, lanes 7-9, and S3c).

**Figure 3.**
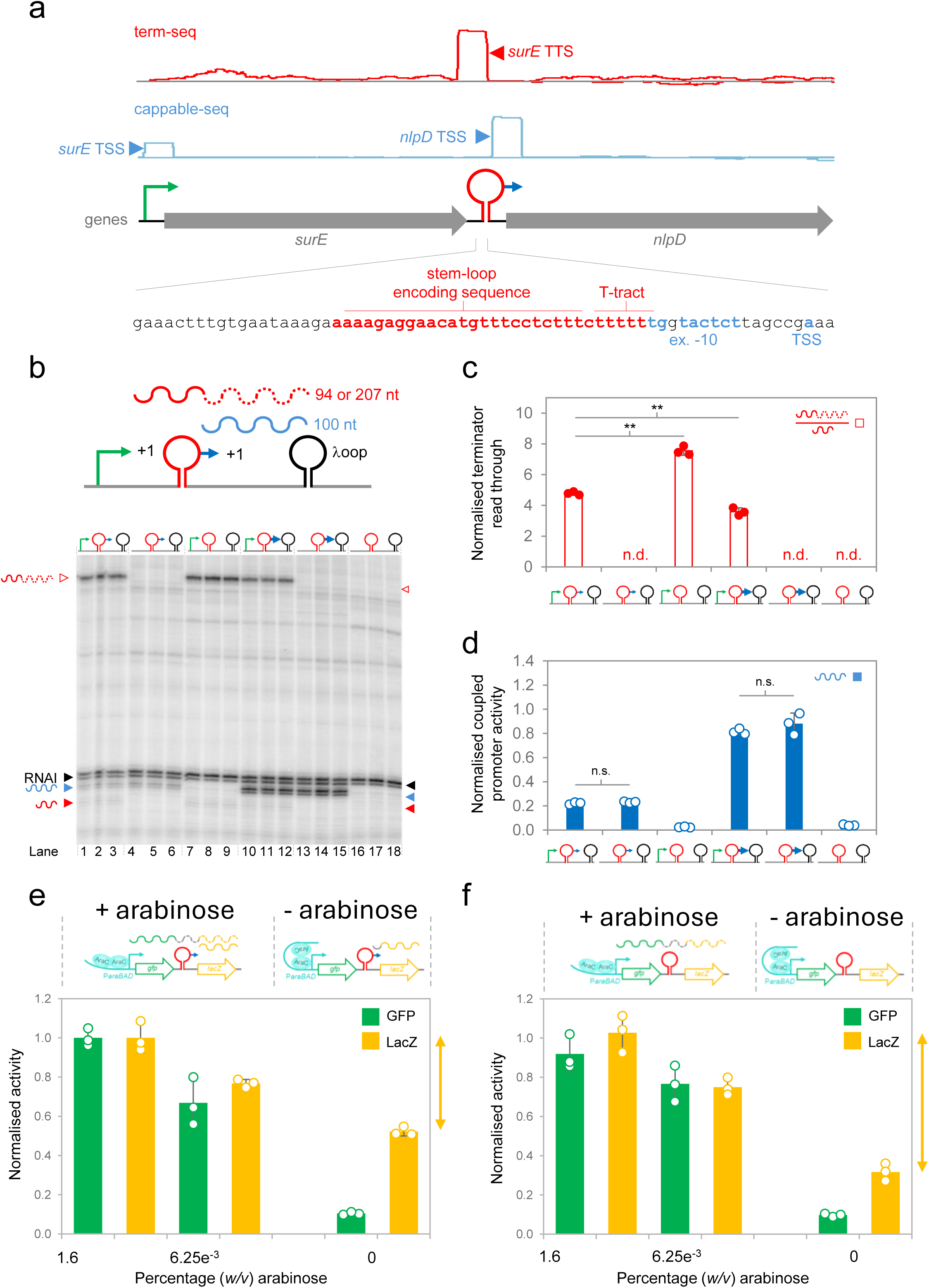
Activity of the *surE* terminator is enhanced by the coupled *nlpD* promoter. **a. The *surE* terminator and *nlpD* promoter are physically coupled.** A schematic of the *surE*-*nlpD* locus is shown alongside data from RNA-seq, term-seq and cappable-seq experiments. The transcription start sites (TSSs) for *surE* and *nlpD* are labelled, as is the site of *surE* mRNA termination. The DNA sequence expansions show base composition of the *surE* promoter (green bent arrow) and the *nlpD* promoter-coupled *surE* terminator (red lollipop and blue overlapping arrow). Key DNA sequence elements are colour coded and labelled. Exact mutations to inactivate the *surE* or *nlpD* promoters (mut1 or mut2 respectively) are shown in Figure S2. **b. DNA template used for *in vitro* transcription assays.** The *surE* terminator (red lollipop) coupled with the *nlpD* promoter (blue arrow) was cloned between the *surE* promoter (green arrow) and λ*oop* terminator. Transcription from the *surE* promoter can terminate at the *surE* terminator, generating a 94 nucleotide (nt) RNA, or read through to λ*oop* so that a 207 nt transcript is made. Transcription from the coupled *nlpD* promoter terminates at λ*oop* to generate a 100 nt RNA. The RNAI transcript serves as an internal control. The DNA templates for *in vitro* transcription are shown schematically above the gel lanes. Reactions were done in triplicate. **c,d. Activity of the *surE* terminator is controlled by the coupled *nlpD* promoter *in vitro*.** The DNA templates for *in vitro* transcription are shown schematically above the gel lanes and are labelled according to the presence of mutations shown in panel b. The RNAI transcript serves as an internal control. Reactions were done in triplicate. The bar charts show relative mean production of the 207 nt read-through *surE* RNA compared to the correctly terminated 94 nt variant (top) and total transcription from the coupled *nlpD* promoter (bottom). Errors bars show standard deviation and overlapping dot plot show data from independent replicates. P was determined using a two-tailed t-test, assuming unequal variance, where n=3. **e,f. Activity of the *surE* terminator is controlled by the coupled *nlpD* promoter *in vivo*.** The *nlpD*-promoter-coupled *surE* terminator was cloned between GFP and *lacZ*. Transcription of the former gene depends on the upstream *araBAD* promoter, whilst transcription of *lacZ* can occur due to *surE* terminator read though or transcription from the *nlpD* promoter. The left hand bar chart shows mean GFP and β-galactosidase activity values with or without different concentrations of arabinose. Expected transcription is indicated by the associated schematic. Increased *lacZ* expression due to arabinose addition (indicated by orange double headed arrow) must be due to terminator read through. The adjacent graph shows equivalent data in the absence of the overlapping *nlpD* promoter. Increased β-galactosidase activity, induced by arabinose, is greater when the *nlpD* promoter is absent. Error bars show standard deviation (n=3) and dots are values for each replicate.

### Coupled promoter activity modulates terminator efficiency in vivo

Next, we cloned the promoter-coupled *surE* terminator between *gfp* and *lacZ*. Here, expression of GFP is controlled by the *araBAD* promoter (Figure 3e, schematic). Hence, in the absence of arabinose, GFP activity is low and the detected β-galactosidase expression must depend on the terminator-coupled *nlpD* promoter (Figure 3e bar chart). Addition of arabinose increased both GFP expression and β-galactosidase activity. The 1.9-fold increase in the latter (indicated by the double headed arrow) must be due to P*araBAD* and terminator read through. In a repeat set of experiments, with the terminator-coupled *nlpD* promoter inactivated, arabinose caused a larger 3.3-fold increase in β-galactosidase activity (Figure 3f). Thus, consistent with our observations *in vitro*, loss of the *nlpD* promoter reduced coupled-terminator efficiency. In similar experiments, with the semi-synthetic promoter-coupled terminator, we inactivated the upstream, or terminator-coupled, promoters. Read-through transcription, revealed by reduced *lacZ* expression when the upstream promoter is mutated, was only evident when the coupled promoter was also absent (Figure S3d).

### Control of termination efficiency by a coupled promoter relies on proximity but not supercoiling in vitro

We reasoned that overlapping promoters might enhance termination in two ways. First, the elongation complex could collide with promoter-bound RNA polymerase and pause; a pre-requisite for termination^13^. Second, positive supercoils, which also induce pausing, are generated ahead of elongation complexes. Diffusion of these supercoils could be trapped by the downstream initiating RNA polymerase^33,34^. Whilst the former model requires promoter-terminator proximity, supercoiling effects should be lost if DNA is linearised. To understand the role of proximity, we inserted from 20-100 bp of DNA between the *surE* terminator and coupled *nlpD* promoter (Figures 4a and S2). As a control, we also mutated each template to remove the upstream *surE* promoter. For each DNA template, we measured read-through transcription and coupled *nlpD* promoter activity (Figures 4b-d). Read through the *surE* terminator increased when uncoupled from the *nlpD* promoter by inserted DNA (Figure 4c). This effect required insertions of at least 35 bp, with larger insertions having little impact. Conversely, *nlpD* promoter activity was unaltered (Figure 4d). As expected, *surE* transcripts were lost upon *surE* promoter mutation (Figure 4b, lanes marked “-“).

**Figure 4.**
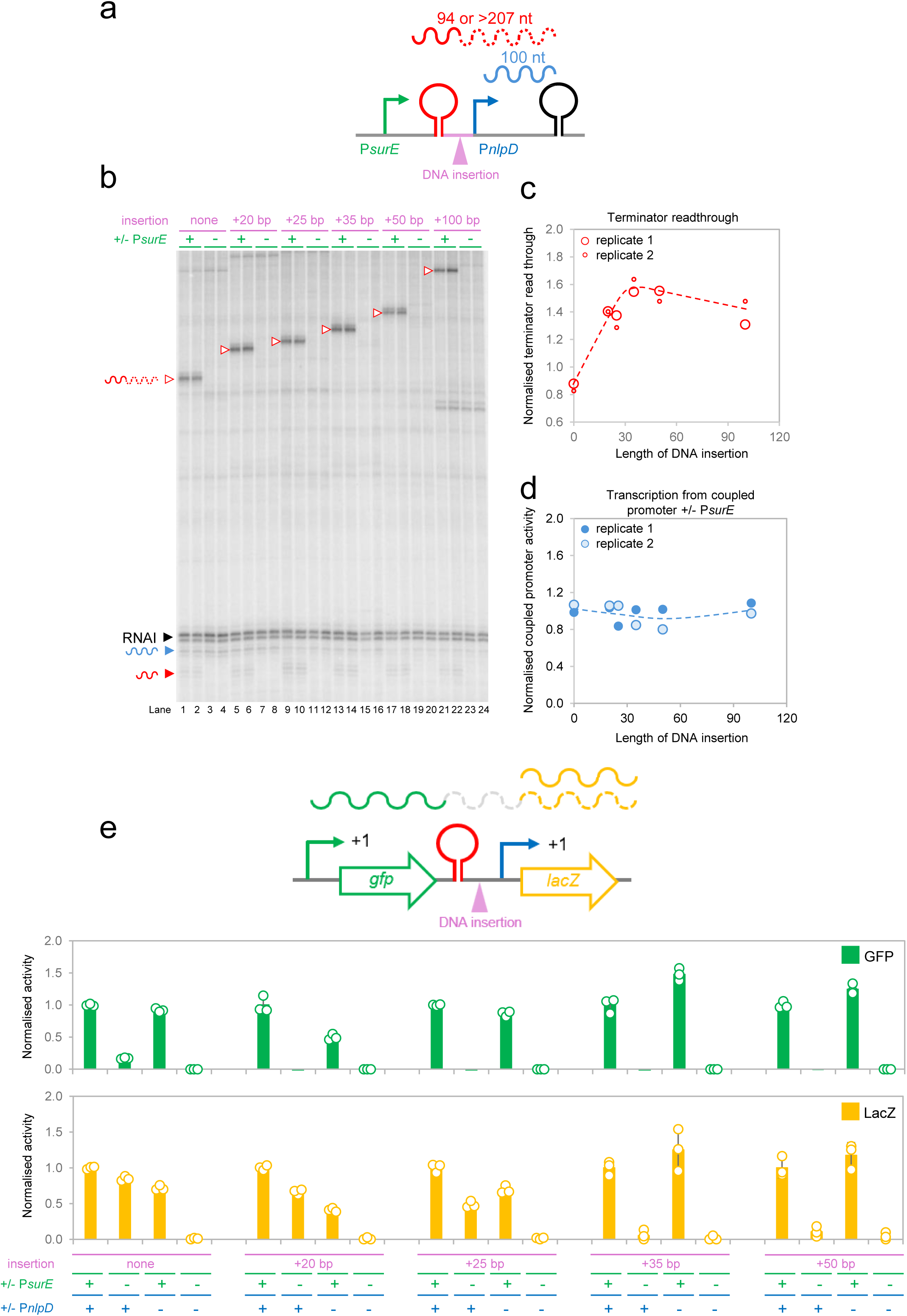
Read-through the *surE* terminator increases upon separation from the coupled *nlpD* promoter. **a. Schematic representation of DNA templates for *in vitro* transcription.** The starting DNA template (top) results consists of the *surE* terminator (red lollipop) coupled to the *nlpD* promoter (blue arrow). This promoter-coupled terminator is positioned between the *surE* promoter (green) and λ*oop* terminator (black). Expected RNA products are labelled by size in nucleotides (nt). Variants that uncouple the *nlpD* promoter from the *surE* terminator (middle) have DNA insertions of between 20 bp and 100 bp (pink). Derivatives lacking the *surE* promoter were also generated (bottom). **b. *In vitro* transcription assays show increased *surE* terminator readthrough upon separation from the *nlpD* promoter.** The gel image shows results of *in vitro* transcription assays using the DNA templates shown in panel a. Reactions were done in duplicate and the RNAI transcript serves as an internal control. **c,d. Quantification of *in vitro* transcription.** The line graphs plot quantifications of *surE* terminator read though or *nlpD* promoter activity. Data from replicates 1 and 2 are plotted as large and small dots respectively. To calculate terminator readthrough, the ratio of bands corresponding to the terminated (94 nt) and read through (207 nt or greater) *surE* transcripts was determined. For transcription from the *nlpD* promoter, we took a similar approach but compared activities +/- the upstream *surE* promoter. **e. Quantification of *surE* terminator read though *in vivo*.** The schematics show different dual-reporter constructs. Genes are shown as block arrows. Otherwise, labelling is as in panel a. The upper and lower bar charts shows mean GFP and β-galactosidase activities respectively. Error bars show standard deviation (n=3) and dots shown replicate values.

To understand the importance of supercoiling, we repeated the analyses shown in Figures 3b and S3, using linearised DNA templates, which allow supercoils to dissipate (Figure S4). Changes in termination efficiency, induced by coupled-promoter mutation, resembled our prior results. Hence, physical proximity of a coupled promoter, but not DNA supercoiling constraints, enhanced termination efficiency.

### Control of termination efficiency by a coupled promoter relies on proximity in vivo

To confirm the impact of proximity *in vivo*, we used the dual *gfp*-*lacZ* reporter system described above. The *surE* promoter was cloned upstream of *gfp*, and the promoter-coupled *surE* terminator between *gfp* and *lacZ* (Figures 4e and S2). As expected, mutating the upstream *surE* promoter abolished GFP activity, whilst changes to the *nlpD* promoter, coupled to the *surE* terminator, had little effect (Figure 4e, green bars). Consistent with our model, *surE* promoter loss had little impact on β-galactosidase activity if the *nlpD* promoter, and coupled terminator, were correctly spaced (Figure 4e, first group of orange bars). Conversely, loss of the *surE* promoter did impact *lacZ* expression if the *nlpD* promoter was sufficiently far from the coupled terminator. The maximum effect, due to inactivating the *surE* promoter, was seen when at least 35 bp of DNA was inserted (Figure 4e, fourth group of orange bars). Taken together, our observations are consistent with coupled promoters aiding termination as a function of TTS proximity.

### Promoter-coupled terminators are found in diverse bacterial species

To understand promoter-terminator coupling in other bacteria, we applied cappable-seq and term-seq to *Escherichia coli* and *Bacillus subtilis*. The data for these species, separated by ∼2 billion years of evolution, are summarised in Figure 5a^35^. We identified 540 and 109 coupled TTS-TSS pairs, separated by ≤ 35 bp, for the two organisms respectively (Table S1). Consistent with our observations above, TTSs occurred most frequently just upstream of promoters, in the same intergenic region, between tandem genes (Figure 5b). Figure 5c shows high-resolution distance frequency plots for all species tested in this work. For each organism, there is a spacing incidence peak for TTS-TSS pairs ∼10 bp apart. As described above for *A. baumannii*, this spacing preference was also evident using *de novo* bioinformatic predictions (Figure S1b). Indeed, the pattern was ubiquitous, and also observed for *Klebsiella pneumoniae*, *Pseudomonas fluorescens*, *Deinococcus radiodurans*, *Staphylococcus aureus*, *Streptococcus pneumoniae*, and *Vibrio cholerae* (Figure S1b). We note that computational predictions unambiguously assign each TSS and TTS to a single position. Conversely, experimental tools often detect several adjacent bases for a single promoter or terminator. Further, no inferences can be made for inactive transcription units. Hence, distance profiles, for experimentally identified TSS-TTS pairs, are noisier (compare Figures 5c and S1).

**Figure 5.**
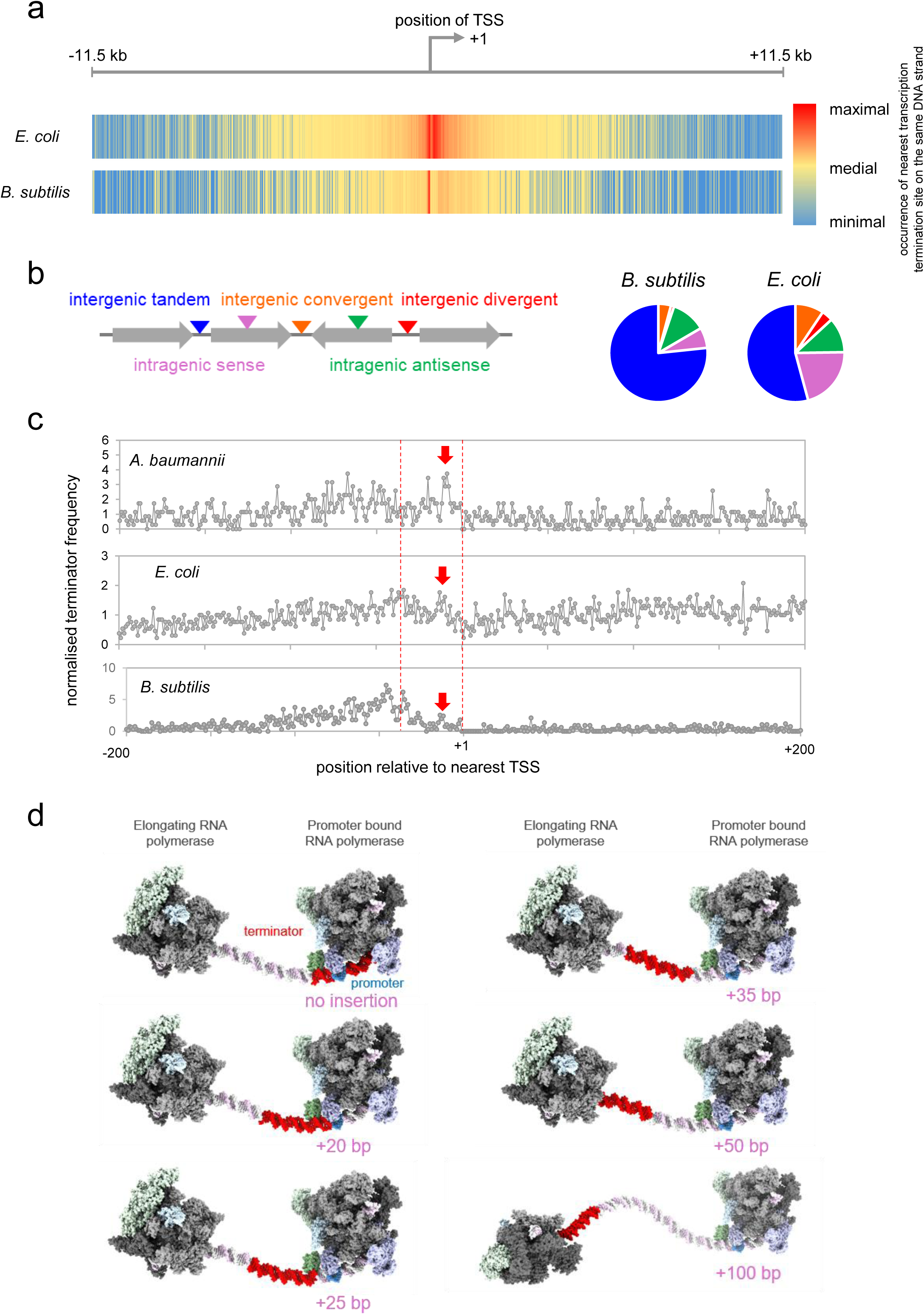
Coupling of promoters to terminators is widespread in distantly related bacteria. **a. Heatmaps showing the global distribution of TTSs with respect to TSSs in *E. coli and B. subtilis*.** All TSSs (from cappable-seq) are aligned at the centre of the heatmap (bent arrow, labelled +1) and colours indicate the abundance of TTSs (from Term-seq) grouped in 100 bp bins, at the relevant position for the same DNA strand. **b. Location of promoter-coupled terminators.** The pie charts show the locations promoter-coupled terminators with respect to genes, in different orientations, as illustrated by the schematic. **c. Conservation of transcription start sites 10 bp downstream of termination sites.** The line graph shows the normalised base pair resolution frequency of transcription termination sites (TTS, y-axis) relative to the position of the nearest transcription start site (TSS, x-axis). The red dashed lines show the 35 bp window within which terminators and promoters overlap. The solid red arrow shows the conserved spacing frequency peak corresponding to ∼10 bp separation. **d. Model for RNA polymerase interference at promoter-coupled terminators.** The RNA polymerase β and β′ subunits are grey. The α N- and C-terminal domains are in pale and dark green respectively (note that the flexible linker between domains was not resolved and one C-terminal domain was detected ^38^). The σ subunit is in lilac. Promoter (blue) and terminator (red) base pairs are highlighted and the size of the insertion between them indicated. The model is based on the promoter-coupled terminator, separating *surE* and *nlpD*, in *A. baumannii*. Protein data bank entries 6PSW^38^ and 7YP9^16^ were used.

### Control of termination by a coupled promoter is common

Our observations suggest a common mechanism used to modulate termination efficiency. To test this, we cloned promoter-coupled terminators, from *E. coli* and *B. subtilis*, between the λ*oop* terminator, and their naturally occurring upstream promoter (Figure S5 schematics). We then quantified upstream and coupled promoter activity, as well as transcriptional readthrough, for each DNA template, and derivatives, using *in vitro* transcription assays. The complete cloned DNA sequences are shown in Figure S2. In all cases, termination efficiency was reduced when the coupled promoter was mutated. The *E. coli gpt* and *ihfA* terminators were extremely efficient and no read-through was detected. Upon coupled promoter mutation, read-through was substantial (Figure S5a and S5b). Although the *B. subtilis yabE* terminator was less efficient, read-through increased >2-fold upon mutation of the coupled *rnmV* promoter (Figure S5c).

### Palindromic sequences encoding terminator stem-loop structures can simultaneously be binding sites for transcription factors

Promoter-coupling results in the terminator stem-loop encoding sequence being directly upstream of, or overlapping, a promoter -35 hexamer. Transcription factors (TFs) bind palindromic DNA elements in such locations. We speculated that, at some promoter-coupled terminators, the sequence encoding the RNA stem-loop might also be recognised by a TF. Comparison of our term-seq analysis for *E. coli*, with TF binding analyses, revealed that the *yobI* terminator palindrome is a LexA binding site for *ruvA* repression (Figure S6a)^36^. Binding of LexA to its target blocks transcription from the *ruvA* promoter and reduces readthrough of the *yobI* terminator. This occurs even if the *ruvA* promoter is inactivated by mutation (Figures S6b and S6c).

## CONCLUSIONS

Bacterial genomes are compact and we show that promoter-terminator coupling optimises use of intergenic space^37^. This is possible because dual-purpose DNA sequences act as both terminator and promoter elements. Indeed, in some cases, such sequences are also regulator binding sites (Figure S6). Hence, multiple control signals can be encoded by the same short DNA region. Conserved in distantly related bacteria (Figures 5 and S1) the primary function of promoter coupling is to regulate termination efficiency (Figures 3 and S3-S5). Mechanistically, this requires that the coupled promoter and terminator overlap (Figure 4). This is consistent with structural models of elongating and promoter bound RNA polymerase (Figure 5d)^16,38^. Hence, if closer than 35 bp, the promoter bound RNA polymerase occludes some or all of the terminator DNA. Whilst head-to-tail collisions are unlikely to evict promoter-bound RNA polymerase from the template, transcription of the termination signal must require forward translocation of the leading RNA polymerase. This can only happen if extensive contacts between the enzyme and promoter are broken. This likely slows the trailing enzyme, or extends the paused state, during hairpin formation. Although not detected here, in some cases, such collisions may facilitate promoter escape. Most likely, the exact outcome will be determined by the kinetics of initiation and termination.

Their ubiquity suggests promoter-coupled terminators emerged in an ancient ancestor. It is tempting to infer that primordial terminators evolved to optimise the impact of random RNA polymerase collisions around TSSs. Proteins able to bind palindromic DNA, encoding RNA hairpins, might then have been selected due to better regulation of termination or initiation. We note that some prokaryotic promoters bind large transcription factor arrays^39,40^. The cumulative impact of such binding may further support termination. Similarly, the role of elongation and termination factors, at promoter-coupled terminators, should be an area of future interest.

In eukaryotes, DNA folding means that promoters and terminators, for the same transcription unit, can exist in close 3-dimensional space^41^. The process, known as gene looping, facilitates RNA polymerase recycling after transcription terminates^42,43^. Although gene loops have been observed in bacteria, it is not known if these enhance transcription^44^. As an alternative, we speculate that promoter-terminator coupling could also support RNA polymerase recycling. Briefly, recent work has shown that, post-termination, RNA polymerase releases transcripts but remains bound to the DNA^45,46^. Subsequently, the enzyme scans downstream sequences for the next available promoter^45,46^. The likelihood that genomes have evolved to optimally position downstream promoters is therefore high, particularly since aberrant post-termination transcription is toxic^47^. Hence, understanding wider consequences of promoter-terminator coupling is now a key goal. We speculate that future insights could have implications beyond the bacteria. For example, Ni *et al*. showed coupling of terminators and promoters, by dual function DNA elements, in *Saccharomyces cerevisiae*^48^. Whilst these sequences were artificial in origin, naturally occurring instances seem likely to exist.

## METHODS

### Strains, plasmids and oligonucleotides

All strains, plasmids and oligonucleotides used are listed in Table S5. Standard procedures for strain and DNA manipulation were used throughout. All bacterial cultures were grown in Lennox broth (LB) medium unless stated otherwise.

### Cappable-seq and Term-seq

For *E. coli*, we used our previous map of TSSs that combined data from cappable-seq and similar TSS mapping tools^49^. For *B. subtllis*, we used our own prior cappable-seq analysis^50^ and the term-seq data of Kosiński *et al*^51^. In all other cases, RNA 5′ or 3′ ends were mapped with cappable-seq or term seq respectively. Experiments were done twice using total RNA isolated from biological replicates. Strains were grown in LB with shaking at 37 °C until mid-log phase. Aliquots of 2 ml were then pelleted and flash frozen in liquid nitrogen. Cappable-seq and term-seq libraries were prepared by Vertis Biotechnologie AG (Germany) followed by sequencing with an Illumina NextSeq 500 system (75 bp read length). Raw data in FASTQ format are available from ArrayExpress (accession number E-MTAB-17359).

### Bioinformatics

Individual sequence reads in FASTQ format were mapped using Bowtie2^52^. The *A. baumannii*, *E. coli* and *B. subtilis* reference genomes used were those assigned Genbank accession no. NZ_CP008706.1, U00096.3, and NC_000964.3 respectively. Resulting BAM files were used to generate wiggle plots using bam2wig.py^53^. To identify TSSs from cappable-seq data, we located chromosomal positions, on each strand, where the read depth increased more than threefold, compared with the previous base, in both experimental replicates^54^. Bioinformatic identification of putative TSSs, based on genome sequence alone, searched for the motif 5′-TAWWWTNNNNNNR-3′ with no mismatches allowed^55^. For term-seq data, we identified genomic loci where the coverage 4 bp upstream, for the same DNA strand, was at least 2-fold higher and a minimum of 20 reads. For each such position, we then selected instances where the read depth 5 bp downstream was over 3-fold lower. These selection criteria almost always identified multiple adjacent chromosomal positions and the most downstream of these was recorded as the TTS for a given replicate. We then compare replicates and selected only those TTSs identified in both, with a tolerance of +/- 5 bp. Of all termination sites we identified in this way, 83 % were separated by a maximum of 2 bp. To identify putative terminators in DNA sequences we used TransTermHP ^56^.

### Protein purification

The *E. coli* RNA polymerase was purchased from New England Biolabs and σ^70^ was purified as described previously^50^. The *B. subtilis* RNA polymerase was purified using a chromosomal β’ C-terminal domain His_6_ tag fusion^50^. Cells were grown to late exponential phase, in shaking LB medium with 1% (*w/v*) glucose, at 37 °C. Cells were resuspended in lysis buffer (50 mM Tris pH 7.9, 300 mM NaCl, 3 mM β Mercaptoethanol, 5% and cOmplete EDTA-free protease inhibit tablet) and lysed. Lysate was clarified by centrifugation and filtration through. Lysates were applied to a His-Trap HP column (GE Healthcare). Unbound protein was removed by washing the column with lysis buffer then lysis buffer with 25 mM imidazole. Bound protein was eluted in a gradient to 200 mM imidazole in lysis buffer. RNA polymerase containing fractions were pooled, diluted 3-fold in HiTrap buffer (40 mM Tris pH 7.9, 1mM EDTA, 5% glycerol) and loaded onto a HiTrap Q column. RNA polymerase was eluted in a gradient to 1 M NaCl. RNA polymerase containing fractions were pooled, concentrated, and made to 50% glycerol for -20 °C storage. The *B. subtilis sigA* gene, cloned in pET-28a, was overexpressed in T7 Express *E. coli* cells (New England Biolabs) grown in LB to exponential phase at 37 °C with shaking. Protein expression was induced for 3 hours with 1 mM IPTG. Overexpressed σ^A^ forms inclusion bodies, which were solubilised in denaturing His-Trap buffer (20 mM Trips pH 7.9, 5% glycerol, 600 mM NaCl, 8 M urea). Cell debris was removed by centrifugation, and solubilised protein was loaded on to a His-Trap column (GE Healthcare). After washing denaturing His-Trap buffer, stepwise increasing concentrations of imidazole were applied and fractions with σ^A^ were pooled and diluted with an equal volume of denaturing buffer (50 mM Tris pH 7.9, 8 M urea, 10% glycerol, 10 mM MgCl_2_, 10 µM ZnCl_2_, 1 mM EDTA, 10 mM DTT) before dialysis overnight in 2 litres of reconstitution buffer (50 mM Tris pH 7.9, 200 mM NaCl, 20% glycerol, 10 mM MgCl_2_, 10 µM ZnCl_2_, 1 mM EDTA, 1 mM DTT). Refolded σ^A^ was diluted 4 times in HiTrap buffer and loaded onto a HiTrap Q column (GE Healthcare). σ^A^ was eluted in HiTrap buffer with a gradient of 50 mM NaCl to 1 M NaCl. Fractions containing σ^A^ were pooled, concentrated, and made to 50% glycerol for -20 °C storage. Purified LexA was a gift from Douglas Browning ^57^.

### In vitro transcription assays

The *E. coli* RNA polymerase was used for experiments with *A. baumannii* and *E. coli* derived DNA templates. For assays with *B. subtillis* DNA sequences, we used the *B. subtilis* RNA polymerase. Linear DNA fragments (70 ng per reaction) or equivalent sequences cloned in plasmid pSR (300 ng per reaction) were used as DNA templates. Each 20 µl reaction contained 20 mM Tris pH 7.9, 40 mM KCl, 10 mM MgCl_2_, 200 μM ATP/GTP/CTP, 10 μM UTP and 2 μCi [α-^32^P] UTP. Reactions were started by addition of 1 µl *E. coli* RNA polymerase σ^70^ holoenzyme (NEB biolabs) or *B. subtilis* σ^A^ holoenzyme (50 nM). After 10 minutes at 37°C 20 μl of formamide stop solution was added. Reactions were run in either triplicate or duplicate and analysed on the same gel. Electrophoresis was done at ∼80 W for ∼1.5 hours in 1 X TBE running buffer. Gels were dried under vacuum and exposed to a phosphorscreen for 24 hours. Images were scanned using the Amersham Typhoon phosphorimager, with resolution set to 50 μm. Bands of interest were manually identified, with their boundaries defined manually using QuantityOne (Bio-rad). Plasmid pSR also encodes the 108 nt RNAI transcript, which serves as an internal control for reactions with this template. Exact details of quantification are provided in relevant figure legends.

### β-galactosidase and GFP activity assays

Derivatives of plasmid pRW50 were used to transform *E. coli* JCB387 and single colonies were used for each biological replicate. After overnight growth, 200 μl of overnight culture was used to inoculate 5 ml of M9supp media and cells incubated until an OD_650_ of between 0.3 and 0.7, with or without arabinose as indicated. Note that the media choice was taken to reduce background fluorescence. For β-galactosidase assays, cells were lysed with 2 drops each of toluene and 1 % (*v/v*) sodium deoxycholate. Next, 100 μl of lysate was used to measure β-galactosidase activity in glass test tubes according to standard protocols. A no lysate reaction was run as a control. For assays of GFP activity, 200 μl of unlysed cells were transferred to a black round-bottomed 96 well plate. This was done twice for each culture and media only measurements were included as a control. We added 1 drop of chloramphenicol (35 μg/ml) to each well. Plates were incubated at 37°C, with shaking at 200 rpm, for an hour to allow for complete folding of GFP. Fluorescence emission was recorded at a 520 nm, following excitation at 485 nm, and was corrected for subculture cell density and background fluorescence.

## ACKNOWLEDGEMENTS

We would like to thank José Penadés, Yuyi Li, and Cora Chmielowska, for critically reading the manuscript ahead of submission, and Steve Busby, Joeseph Wade and Ramesh Wigneshweraraj for helpful conversations. This work was funded by Leverhulme Trust project grant RPG-2022-191 awarded to DCG and a BBSRC MIBTP studentship awarded to AGF.

## SUPPLEMENTARY FIGURE

**Figure S1.**
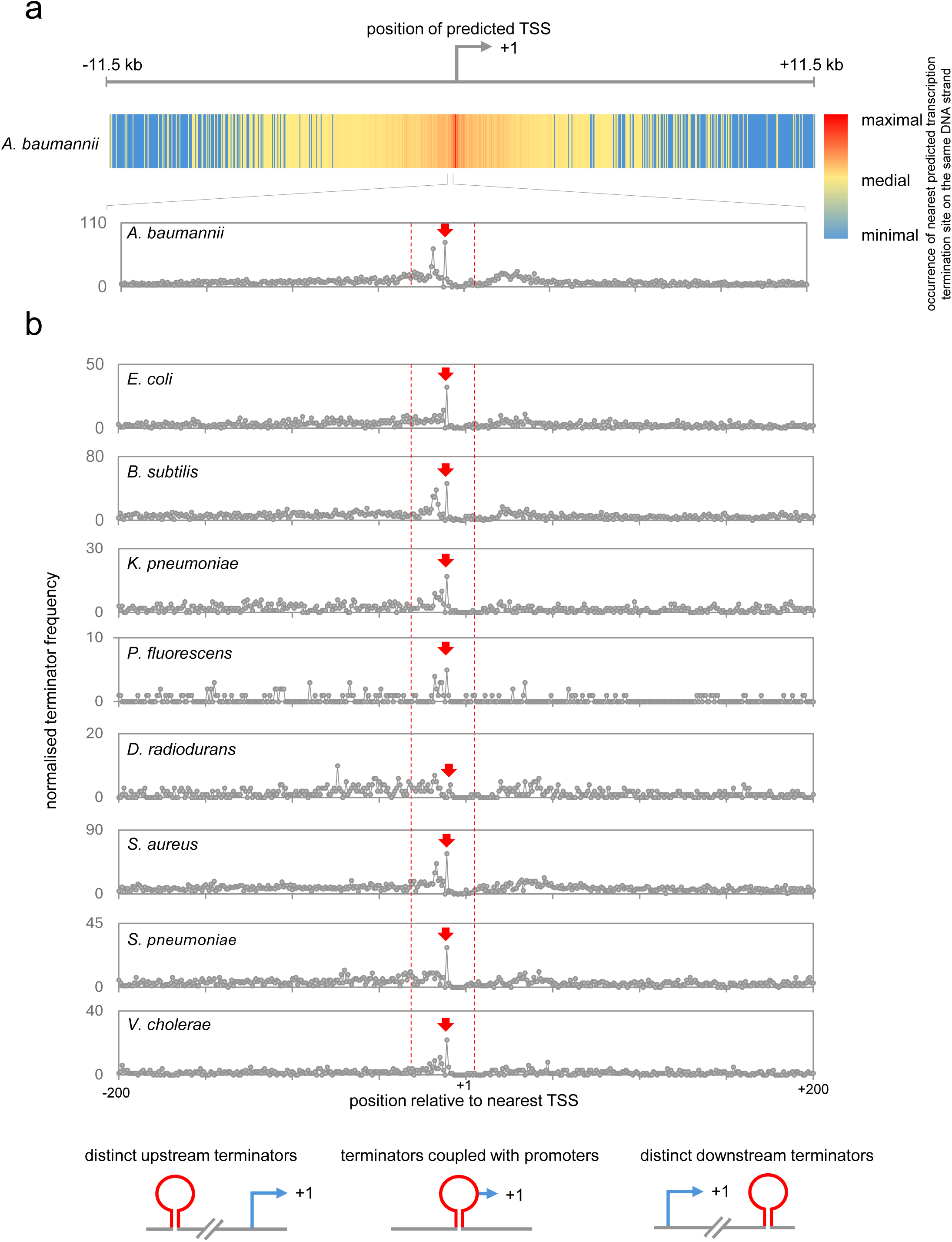
Sequence based prediction of terminator and promoter location identifies promoter-coupled terminators. **a. Heatmap showing global distribution of predicted TTSs and TSSs in *Acinetobacter baumannii*.** All predicted TSSs are aligned at the centre of the heatmap (bent arrow, labelled +1) and colours indicate the abundance of predicted TTSs, grouped in 100 bp bins, at the relevant position for the same DNA strand. The red dashed lines show the 35 bp window within which terminators and promoters overlap. **b.** The line graph shows base pair resolution depiction, of the data in panel a, for *A. baumannii* (top chart) and equivalent data for other bacterial species (lower charts). The red dashed lines show the window within which terminators and promoters overlap. The red block arrows shows the conserved peak for separation of TTS and TSS by ∼10 bp. Note that for some species (e.g. *P. fluorescens*) the number of predicted TSSs and TTSs was much lower, likely because of a higher GC % genome, and so far fewer nearby TTS-TSS pairs were found.

**Figure S2.**
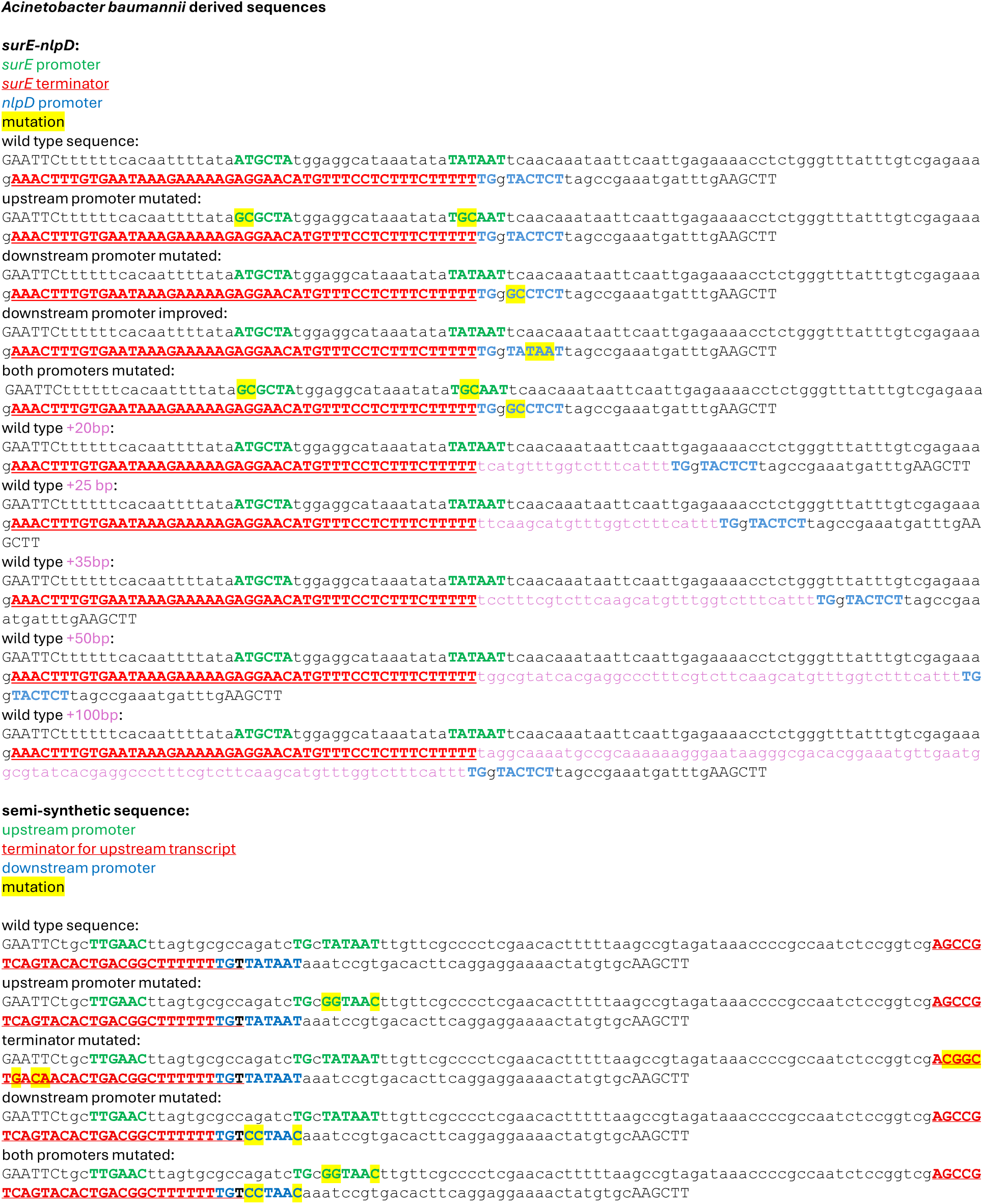
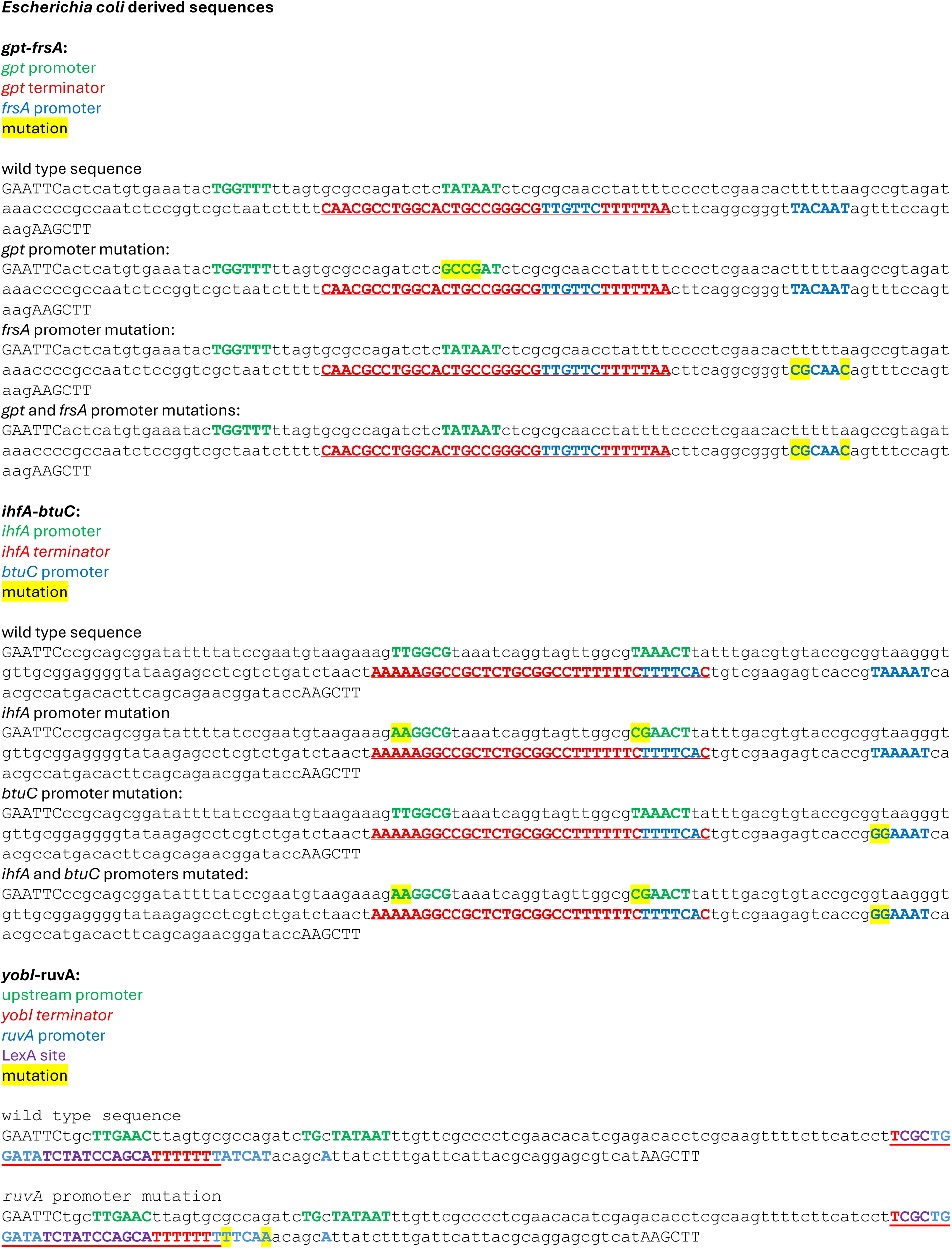
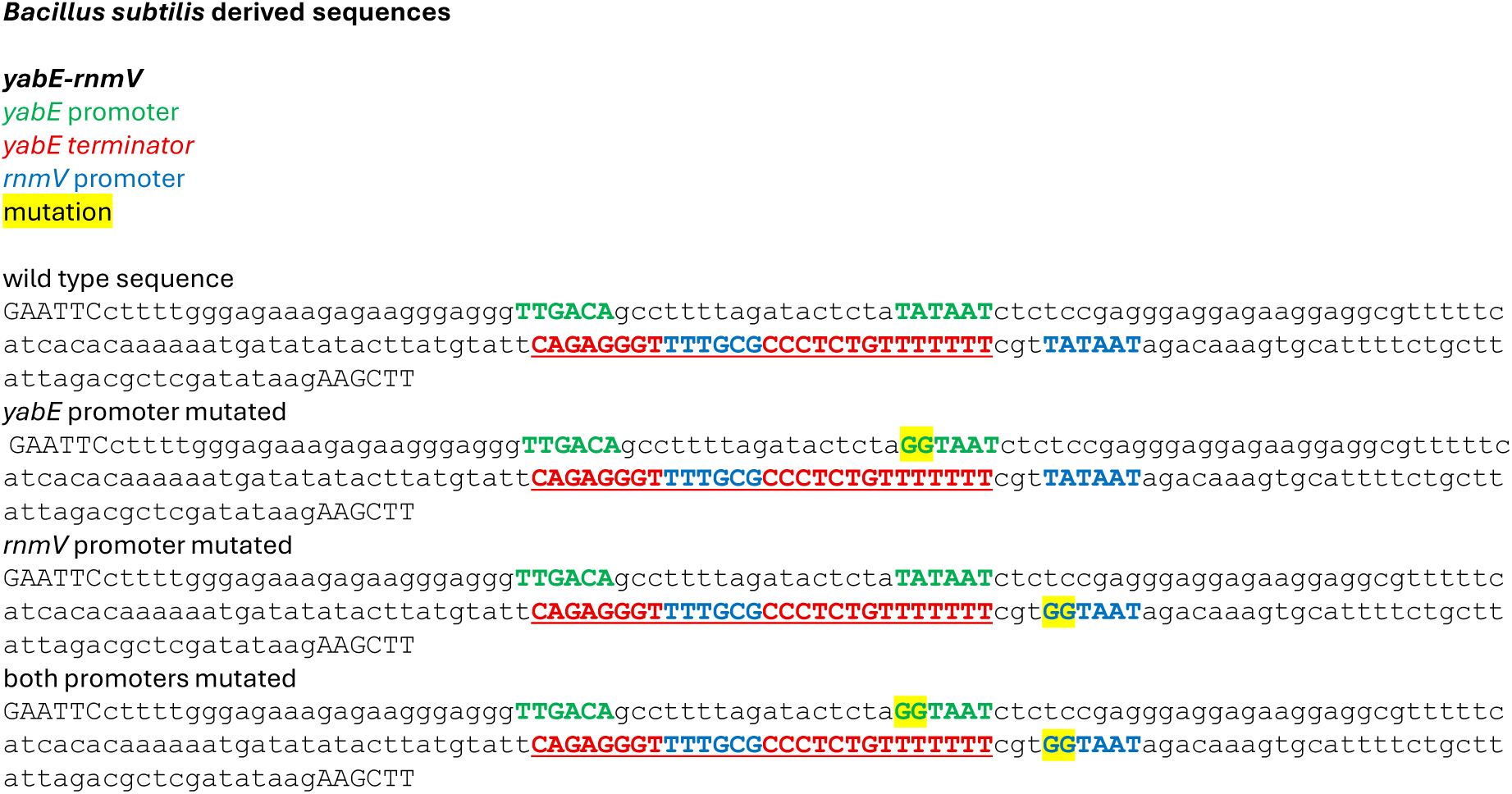
Complete sequences of DNA sections used in this work. Sequences encoding terminators are red. Promoters upstream and coupled to terminators are coloured green and blue respectively. Mutations are highlighted yellow and inserted DNA is pink.

**Figure S3.**
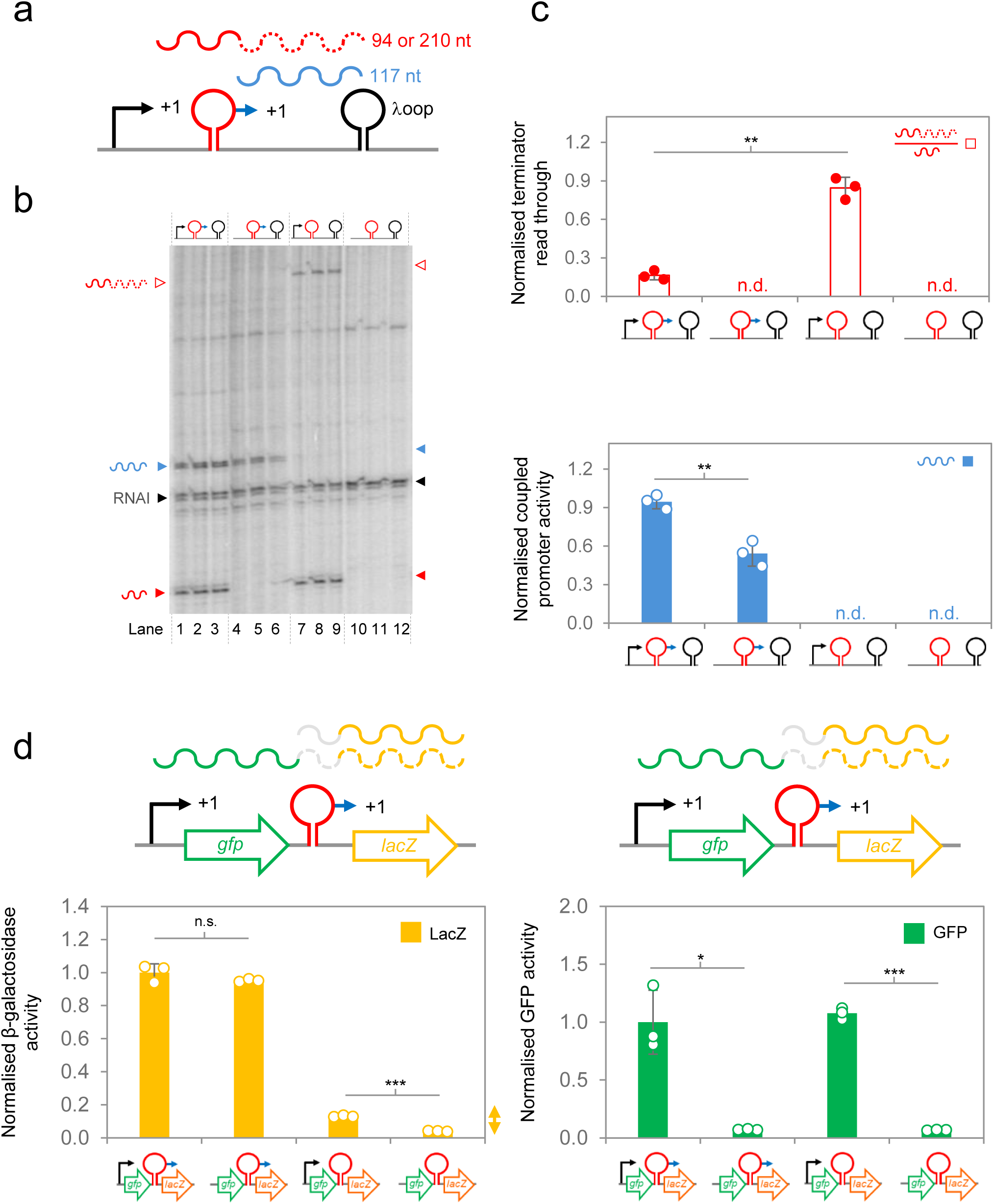
Activity of a semi-synthetic terminator is enhanced by a coupled promoter. **a. Semi-synthetic DNA sequences used to test promoter-terminator coupling.** The terminator (red lollipop) and coupled promoter (blue arrow) were cloned between a constitutive upstream promoter (black arrow) and the λ*oop* terminator (black lollipop). Full sequences are in Figure S2. Transcription from the upstream constitutive promoter can terminate at the promoter-coupled terminator, to generate a 94 nucleotide (nt) RNA, or read through this sequence and make a 210 nt RNA. Transcription from terminator-coupled promoter generates 117 nt RNA. **b. Efficient termination requires a coupled promoter *in vitro*.** The DNA templates for *in vitro* transcription are shown schematically above the gel lanes. Complete sequences are in Figure S2. Note that reactions were done in triplicate. Transcripts are labelled as in panel a and the RNAI transcript serves as an internal control. **c. Quantification of terminator read through and coupled promoter activity.** The bar charts show relative levels of transcriptional read through (top) and total transcription from the coupled promoter (bottom). Errors bars show standard deviation (n=3) and overlapping dot plots show data from independent replicates. Read through was calculating using the ratio of terminated (94 nt) to read through (210 nt) transcription. For transcription from the coupled promoter, band intensity was normalised using the RNAI signal. Significance was calculated using a paired t-test assuming unequal variance. **d. Efficient termination requires a coupled promoter *in vivo*.** The gene encoding GFP was cloned between the constitutive upstream promoter, and promoter coupled terminator, shown in panel a. Expression of the downstream *lacZ* gene is dependent on terminator read through and/or the coupled promoter. The bar charts show mean β-galactosidase and GFP activity values. Error bars show standard deviation (n=3) and the overlayed dot plot shows data from each replicate. Changes in β-galactosidase activity, when the upstream constitutive promoter (black arrow) is deleted, must be due to terminator read through. Such changes only occur if the promoter coupled terminator (blue arrow) is absent. Significance was calculated using a paired t-test assuming unequal variance.

**Figure S4.**
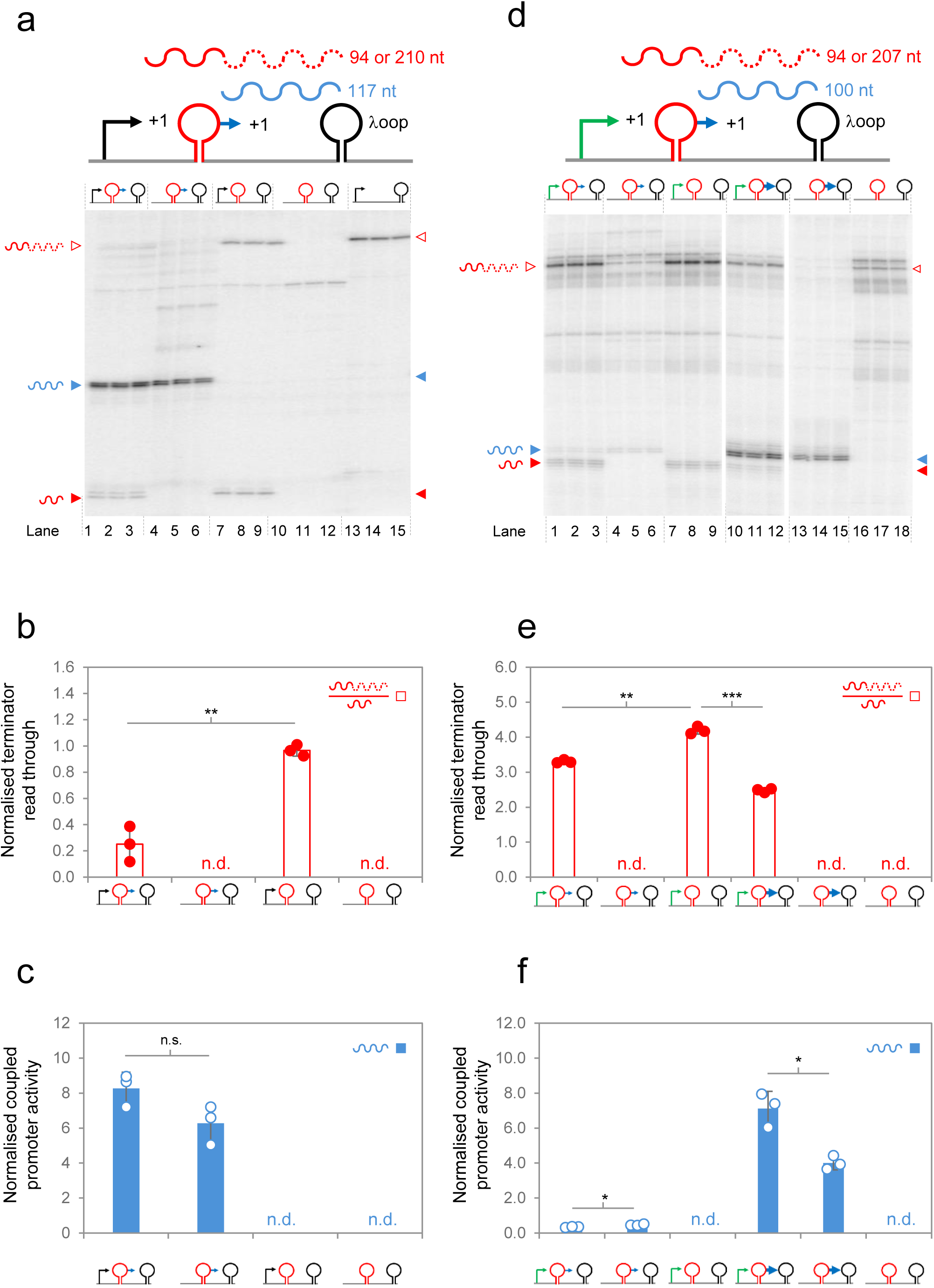
Control of termination efficiency by coupled promoters does not require DNA supercoiling. **a,d. Termination efficiency is controlled by a coupled promoter in the absence of DNA supercoiling.** The DNA templates for *in vitro* transcription, and their variants, are shown schematically above the gel images. Reactions were done in triplicate. **b-f. Quantification of terminator read through and coupled promoter activity.** The bar charts show relative levels of transcriptional read through (red) and total transcription from the coupled promoter (blue). Errors bars show standard deviation (n=3) and overlapping dot plots show data from independent replicates. Read through was calculated using the ratio of terminated to read through transcription. For transcription from the coupled promoter, band intensity was normalised using the RNAI signal. Significance was calculated using a paired t-test assuming unequal variance.

**Figure S5.**
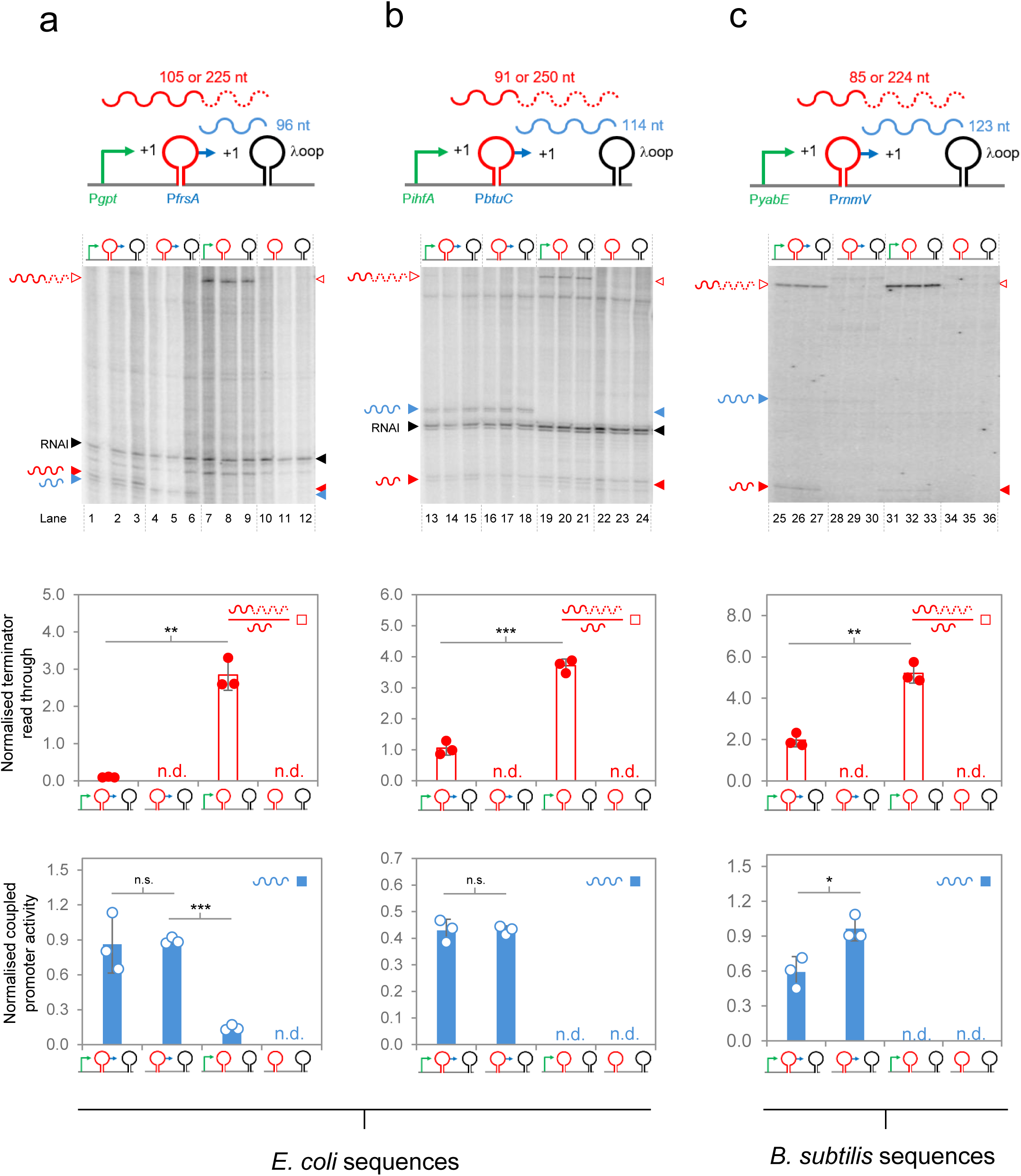
Properties of promoter-coupled terminators isolated from *Escherichia coli* and *Bacillus subtilis*. **a,b,c. DNA templates and variants used for *in vitro* transcription assays.** The DNA templates for *in vitro* transcription, and their variants, are shown schematically above the gel images. **d-g. Quantification of terminator read through and coupled promoter activity.** The bar charts show relative levels of transcriptional read through (red) and total transcription from the coupled promoter (blue). Errors bars show standard deviation (n=3) and overlapping dot plots show data from independent replicates. Read through was calculating using the ratio of terminated to read through transcription. For transcription from the coupled promoter, band intensity was normalised using the RNAI signal, except for the *B. subtilis* example, which used a linear DNA template, where we used the raw band intensities. Significance was calculated using a paired t-test assuming unequal variance.

**Figure S6.**
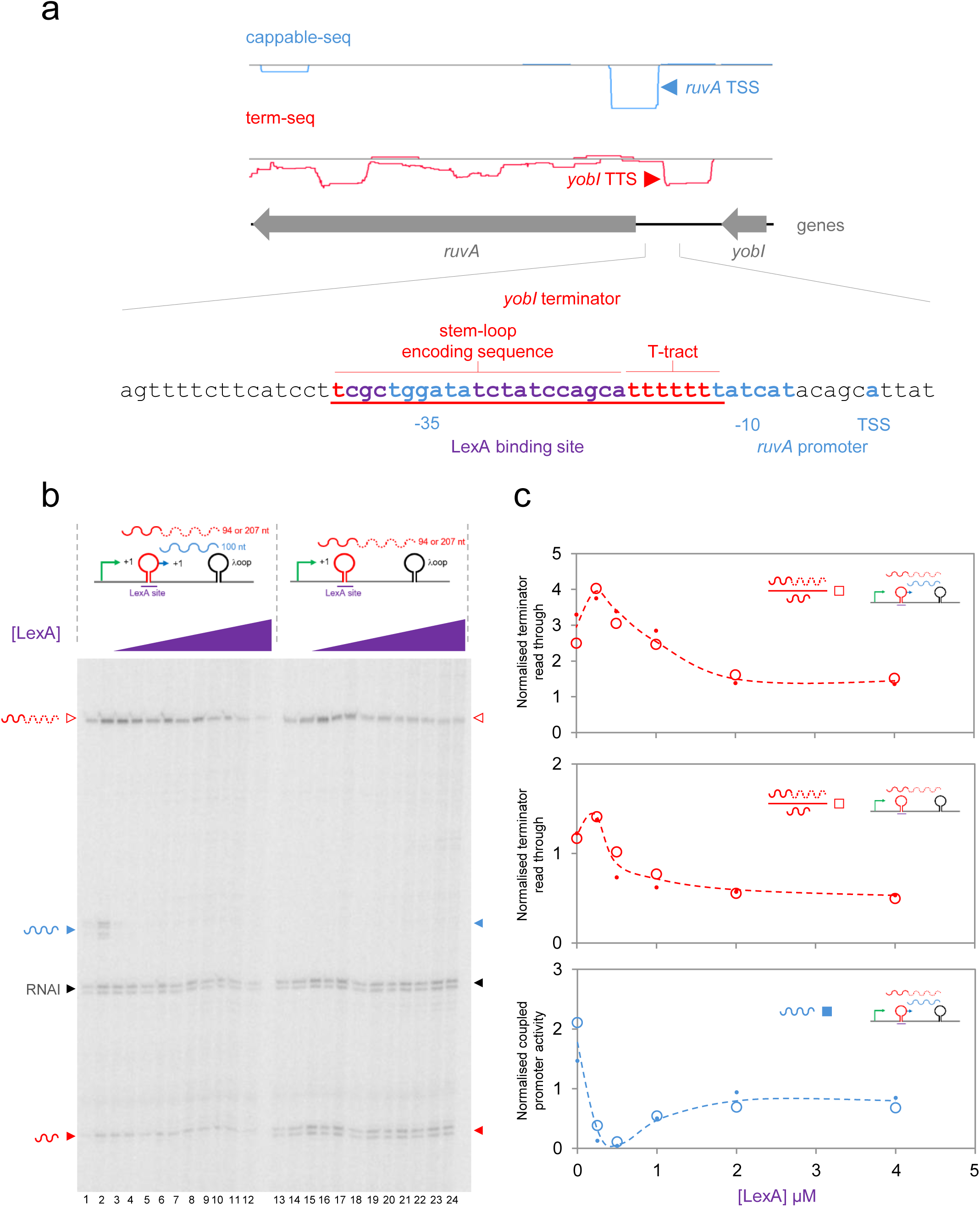
LexA enhances termination of *yobI* transcription by binding a site precisely overlapping the terminator stem-loop DNA motif. **a. Sequence of the *E. coli yobI*-*ruvA* intergenic DNA.** Key DNA sequences are labelled. We note that Shinagawa *et al*. previously proposed two *ruvA* transcription start sites (indicated by grey dots)^58^. We did not detect any RNA 5′ ends aligned with these loci and the associated promoter elements are a poor match to the consensus for σ^70^ binding. **b. LexA enhances termination of the *yobI* promoter.** The gel image shows results of *in vitro* transcription experiments. Each reaction was done in duplicate and replicates are in consecutive lanes. **c. Quantification of terminator read through and coupled promoter activity.** The bar charts show relative levels of transcriptional read through (red) and total transcription from the coupled promoter (blue). Errors bars show standard deviation (n=3) and overlapping dot plots show data from independent replicates. Read through was calculating using the ratio of terminated to read through transcription. For transcription from the coupled promoter, band intensity was normalised using the RNAI signal.

**Figure S7.**
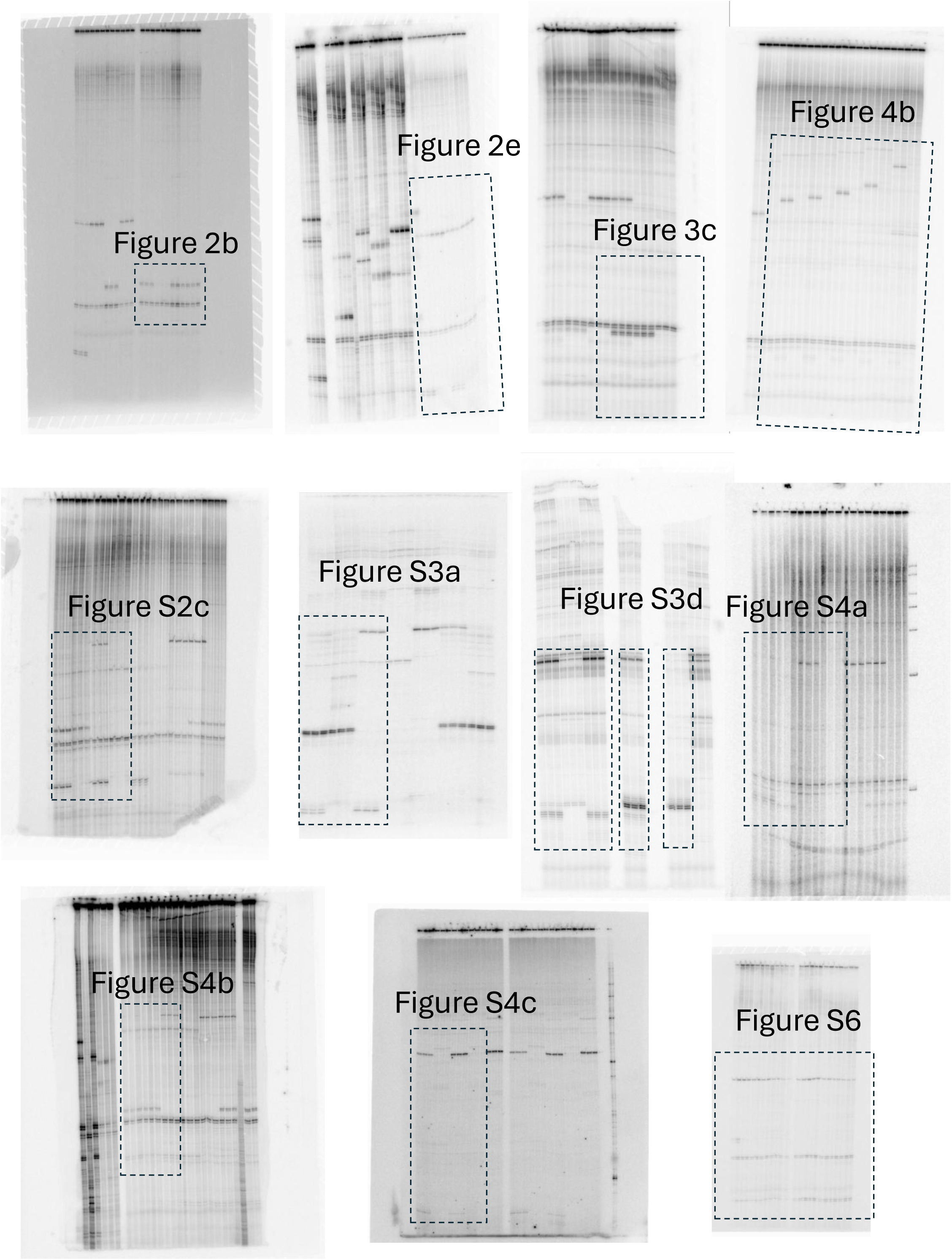
Raw gel images.

