## Supplementary material for "Widespread coupling of promoters and terminators of transcription in bacteria": Table S5

Table S5: Strains, oligonucleotides and synthetic DNA fragments

*Bacterial strains*

*Name	Description	Source*

*A. baumannii* AB5075	Highly virulent and drug resistant isolate from an osteomyelitis	^1^

tibial infection.

*E. coli* JCB387	Used for general plasmid DNA manipulation. ∆*nirB*∆*lac*.	^2^

*E. coli* MG1655	K-12 F^-^ λ^-^ *ilvG*^-^ *rfb*-50 *rph*-1	^3^

*E. coli* DH5α	Used for general plasmid DNA manipulation. *fhuA*2Δ(*argF*-*lacZ*)U169	NEB

*phoA* *glnV*44 Φ80Δ(*lacZ*)M15 gyrA96 *recA*1 *relA*1 *endA*1 *thi*-1 *hsdR*17

*E. coli* BW25113	F⁻ Δ(*araD*-*araB*)567, Δ*lacZ*4787(::*rrnB*-3), λ⁻, rph-1, Δ(*rhaD*-*rhaB*)568,	^4^

*hsdR*514, *rph*-1

*B. subtilis* 168ca	*trpC*2	^5^

*Plasmids*

*Name	Description	Source*

pRW50	low copy number *lac* fusion vector with *Eco*RI/*Hin*dIII cloning site. *oriV* origin.	^6^

TetR.

pSR	pBR322-derived plasmid. Features cloning site upstream of a λoop	^7^

transcription terminator. AmpR.

pBAD30	Vector for tightly controlled, L-arabinose-inducible protein expression.	^8^

Uses the *araBAD* promoter and the AraC regulator. Used as a template

to generate a DNA fragment to be cloned upstream of *gfp* in pRW50.

p15A origin. Amp^R^.

pVRL2::*gfp*	Medium copy number plasmid encoding GFP under the control of the *araBAD*	^9^

promoter. Used as a source of the *gfp* gene. ColE1-like origin. Gm^R^.

*Synthetic DNA fragments*^1^

*Name	Sequence	Source*

Semi-synthetic promoter	cttcaa*gaattc*tgcttgaacttagtgcgccagatctgctataatttgttcgcccctcgaacactttttaagccg	Thermo-

-coupled terminator	tagataaaccccgccaatctccggtcgagccgtcagtacactgacggctttttttgttataataaatccgtga	Fisher

cacttcaggaggaaaactatgtgc*aagctt*actccc

*yabE*-*rnmV* promoter-	atcacgaggccctttcgtcttcaa*gaattc*cttttgggagaaagagaagggagggttgacagccttttagat	Genewiz

coupled terminator	actctatataatctctccgagggaggagaaggaggcgtttttcatcacacaaaaaatgatatatacttatgt

attcagagggttttgcgccctctgtttttttcgttataatagacaaagtgcattttctgcttattagacgctcgatat

aag*aagctt*actccccatcccctccagtaatga

*PCR primers for plasmid sequencing*

*Name	Template	Use	Sequence*

pSR F	pSR backbone	Sequencing	gcatttatcagggttattgtctc

pSR R	pSR backbone	Sequencing	catcaccgaaacgcgcgagg

pRW50F	pRW50 backbone	Sequencing	gttctcgcaaggacgagaatttc

pRW50R	pRW50 backbone	Sequencing	aatcttcacgcttgagatac

pBAD30F	pBAD30 backbone Sequencing	atgccatagcatttttatcc

pBAD30R	pBAD30 backbone Sequencing	gatttaatctgtatcagg

pVRL2 F	pVRL2 backbone	Sequencing	actgtttctccatacccgtt

pVRL2 R	pVRL2 backbone	Sequencing	tgggtaacgccagggttttc

*PCR primers for construction of DNA fragments illustrated in Figure 2b*^1,2^

*Name	Template	Use	Cloning strategy	Sequence*

coupled region F	Figure S3a DNA	Synthesising fragment	Digestion-ligation	cttcaa*gaattc*gccaatctccggtcgagc

in Figure 2b

coupled region R	Figure S3a DNA	Synthesising fragment	Digestion-ligation	gggagt*aagctt*gcacatagttttcctcctgaa

in Figure 2b

coupled region	wild type	T-tract mutation	Digestion-ligation	cttcaa*gaattc*gccaatctccggtcgagccg

KO T-tract F	tcagtacactgacggc**cgcgcgc**gttataat

aaatccgtgaca

coupled region	wild type	-35 mutation	Digestion-ligation	cttcaa*gaattc*gccaatctccggtcgagccg

KO-35 element F	tcagt**tgt**ctgacggctttttttgttat

coupled region	wild type	hairpin mutation	Digestion-ligation	cttcaa*gaattc*gccaatctccggtcgagccg

KO hairpin F	tcagtacac**actg**ggctttttttgttataataa

*PCR primers for construction of DNA fragments illustrated in Figure 2e*^1,2^

*Name	Template	Use	Cloning strategy	Sequence*

up region F	Figure S3a DNA	Synthesising fragment	Digestion-ligation	cttcaa*gaattc*tgcttgaacttagtgcgccag

in Figure 2e

up region R	Figure S3a DNA	Synthesising fragment	Digestion-ligation	gggagt*aagctt*acaaaaaaagccgtca

in Figure 2e	gtgtac

up region KO	wild type	T-tract mutation	Digestion-ligation	aggggatggggagt*aagctt*ac**gcgcgcg**

T-tract R	gccgtcagtgtactgacggct

up region KO	wild type	-35 mutation	Digestion-ligation	aggggatggggagt*aagctt*acaaaaaaagc

-35 element R	cgtcag**aca**actgacggctcgaccggaga

up region KO	wild type	hairpin mutation	Digestion-ligation	gggagt*aagctt*acaaaaaaagcc**cagt**gtgt

hairpin R	actgacggctcgaccgg

*Primers used to generate DNA fragment illustrated in Figure S3a*^1,2^

*Name	Template	Use	Cloning strategy	Sequence*

Synthetic F	synthesised DNA	Synthesising fragment	Digestion-ligation	cttcaa*gaattc*tgcttgaacttagtgcgccag	in Figure S3a

Synthetic R	synthesised DNA	Synthesising fragment	Digestion-ligation	gggagt*aagctt*gcacatagttttcctcctgaa	in Figure S3a

KO upstream	synthesised DNA	inactivate upstream	Digestion-ligation	cttcaa*gaattc*tgcttgaacttagtgcgccag

promoter F	promoter	atctgc**gg**taa**c**ttgttcgcccctcgaacact

KO coupled	synthesised DNA	inactivate coupled	Digestion-ligation	gggagt*aagctt*gcacatagttttcctcctgaa

promoter R	promoter	gtgtcacggattt**g**tta**gg**acaaaaaaagccg

tcagtgtact

KO terminator R	synthesised DNA	inactivate terminator	Digestion-ligation	gggagt*aagctt*gcacatagttttcctcctgaa

gtgtcacggatttattataacaaaaaaagccg

tcagtgt**tg**t**c**agccgtcgaccggagattggcggggt

*PCR primers for construction of DNA fragment illustrated in Figure S3d*^1,2^

*Name	Template	Use	Cloning strategy	Sequence*

GFP synthetic F	pVRL2::gfp	synthesis of *gfp* encoding	HiFi DNA Assembly	tttttaagccattaaagaggagaaattaagc

section

GFP synthetic R	pVRL2::gfp	synthesis of *gfp* encoding	HiFi DNA Assembly	gtttatctacagcttatttgtatagttcatc

section

synthetic up	FigureS3a	synthesising upstream	HiFi DNA Assembly	ccttgagtccacgctagatct*gaattc*t

GFP F	construct	constitutive promoter	gcttgaacttagtgcgccaga

section

synthetic up	FigureS3a	synthesising upstream	HiFi DNA Assembly	tcctctttaatggcttaaaaagtgttcga

GFP R	construct	constitutive promoter	gggg

section

synthetic	FigureS3a	synthesising semi-synthetic	HiFi DNA Assembly	caaataagctgtagataaaccccgccaatct

coupled GFP F	construct	promoter-coupled terminator

section

synthetic	FigureS3a	synthesising semi-synthetic	HiFi DNA Assembly	tctttttgcgcactgacaa*aagctt*gcacatag

coupled GFP R	construct	promoter-coupled terminator	ttttcctcctgaag

section

*Primers used to generate the gpt-frsA promoter-coupled terminator (Figure S5a)*^1,2^

*Name	Template	Use	Cloning strategy	Sequence*

*gpt* F:	*E. coli* genomic	synthesising *gpt-frsA*	HiFi assembly	gtccacgctagatct*gaattc*ACTCATGTGAA

DNA	promoter-coupled terminator	ATACTGGTT

*gpt* R:	*E. coli* genomic	synthesising *gpt-frsA*	HiFi assembly	GATTGGCGGGGTTTATCTACGGCTT

DNA	promoter-coupled terminator	AAAAA

*frsA* F:	*E. coli* genomic	synthesising *gpt-frsA*	HiFi assembly	GTAGATAAACCCCGCCAATCTCCGG

DNA	promoter-coupled terminator	TCGCT

*frsA* R:	*E. coli* genomic	synthesising *gpt-frsA*	HiFi assembly	tttgcgcactgacaa*aagctt*CTTACTGGAAA

DNA	promoter-coupled terminator	CTATTGTAAC

KO *gpt* F:	wild type	mutating *gpt* promoter	HiFi assembly	agatct*gaattc*ACTCATGTGAAATACTGG

TTTTTAGTGCGCCAGATCTCGCCGA

TCTC**GCGC**AACCTATTTTCCCCT

KO *frsA* R:	wild type	mutation *frsA* promoter	HiFi assembly	tgacaa*aagctt*CTTACTGGAAACT**G**TTG

**CG**ACCCGCCTGAAGTTAAAAAG

*Primers used to generate the ihfA-btu promoter-coupled terminator (Figure S5b)*^1,2^

*Name	Sequence	Cloning strategy	Sequence*

*ihfA F*:	*E. coli* genomic	synthesising *ihfA-btuC*	HiFi assembly	gtccacgctagatct*gaattc*CCGCAGCGGATA

DNA	*promoter-coupled terminator*	TTTTATCC

*ihfA* R:	*E. coli* genomic	synthesising *ihfA-btuC*	HiFi assembly	TTAGATCAGACGAGGCTCTTATACCC	DNA	*promoter-coupled terminator*	CTCC

*btuC* F:	*E. coli* genomic	synthesising *ihfA-btuC*	HiFi assembly	AAGAGCCTCGTCTGATCTAACTAAA	DNA	*promoter-coupled terminator*	AAGGC

*btuC* R:	*E. coli* genomic	synthesising *ihfA-btuC*	HiFi assembly	tttgcgcactgacaa*aagctt*GGTATCCGTTCT

DNA	*promoter-coupled terminator*	GCTGAAGTG

KO *ihfA* F:	wild type	mutating *ihfA* promoter	HiFi assembly	agatct*gaattc*CCGCAGCGGATATTTTATC

CGAATGTAAGAAAG**AA**GGCGTAAATC

AGGTAGTTGGCG**CG**AACTTATTTGACGTGTACCGCGGT

KO *btuC* R:	wild type	mutating *btuC* promoter	HiFi assembly	tgacaa*aagctt*GGTATCCGTTCTGCTGAA

GTGTCATGGCGTTGATTT**CC**CGGTGACTCTTCGACAG

*Primers used to generate the yobI-ruvA promoter-coupled terminator (Figure S6)*^1,2^

*Name	Sequence	Cloning strategy*

Upstream DNA F:	*E. coli* genomic	synthesising upstream	HiFi assembly	CTTCAA*GAATTC*tgcTTGAACttagtgcgc

DNA	promoter region	cag

Upstream DNA R:	*E. coli* genomic	synthesising upstream	HiFi assembly	gtgtctcgatgtgttcgaggggcgaacaaA

DNA	promoter region

*ruvA* F:	*E. coli* genomic	*yobI* terminator coupled	HiFi assembly	cctcgaacacatcgagacacctcgcaagtt

DNA	to *ruvA* promoter

*ruvA* R:	*E. coli* genomic	*yobI* terminator coupled	HiFi assembly	GAGT*AAGCTT*atgacgctcctgcgtaatga

DNA	to *ruvA* promoter

KO *ruvA* R:	*E. coli* genomic	*ruvA* promoter mutation	HiFi assembly	GAGT*AAGCTT*atgacgctcctgcgtaatgaat

DNA	caaagataatgctgt**t**tga**a**aaaaaaatgctgga

tagatat

Primers used to generate fragment with the *surE* promoter, fused to *gfp*, and with the *surE* terminator and *nlpD* promoter downstream, for cloning in pRW50 to make a *lacZ* fusion in Figure 4e^1,2,3^

*Name	Template	Use	Cloning strategy	Sequence*

GFP *surE* F	pVRL2::gfp	GFP section	HiFi Assembly	tcgagaaagaattaaagaggagaaattaag

GFP *surE* R	pVRL2::gfp	GFP section	HiFi Assembly	tcacaaagttagcttatttgtatagttcat

*surE* GFP F	genomic DNA	*surE* promoter section	HiFi Assembly	tccacgctagatct*gaattc*ttttttcacaattttataatg

*surE* GFP R	genomic DNA	*surE* promoter section	HiFi Assembly	cctctttaattctttctcgacaaataaacc

*nlpD* GFP F	genomic DNA	*nlpD* promoter section	HiFi Assembly	caaataagctaactttgtgaataaagaaaa

*nlpD* GFP R	genomic DNA	*nlpD* promoter section	HiFi Assembly	ttgcgcactgacaa*aagcttc*aaatcatttcggctaa

gagtaccaaaaaag**a**aagaggaaacatgttcc

Primers used to insert DNA between the *surE* terminator, and *nlpD* promoter, in derivatives of pSR or pRW50^1^

*Name	Template	Use	Cloning strategy	Sequence*

*surE* + 20bp F	wild type	Inserting 20 bp	Q5 mutagenesis	ctttcattttggtactcttagccgaaatgatt

*surE* + 20bp R	wild type	Inserting 20 bp	Q5 mutagenesis	accaaacatgaaaaaagaaagaggaaacatgt

*surE* + 25bp F	wild type	Inserting 25 bp	Q5 mutagenesis	tcaagcatgtttggtctttcattttggtactc

*surE* + 25bp R	wild type	Inserting 25 bp	Q5 mutagenesis	aaaaaag**a**aagaggaaacatgttcctcttttt

*surE* + 35bp F	wild type	Inserting 35 bp	Q5 mutagenesis	cgtcttcaagcatgtttggtctttcattttggtac

*surE* + 35bp R	wild type	Inserting 35 bp	Q5 mutagenesis	aaaggaaaaaag**a**aagaggaaacatgttcctcttt

*surE* + 50bp F	wild type	Inserting 50 bp	Q5 mutagenesis	tcacgaggccctttcgtcttcaagcatgtttggtctttca

*surE* + 50bp R	wild type	Inserting 50 bp	Q5 mutagenesis	tacgccaaaaaag**a**aagaggaaacatgttcctcttt

ttct

*surE* + 100bp F	wild type	Inserting 100 bp	Q5 mutagenesis	taagggcgacacggaaatgttgaatggcgtatcacg

aggccctttcgtct

*surE* + 100bp R	wild type	Inserting 100 bp	Q5 mutagenesis	ttcccttttttgcggcattttgcctaaaaaagaaagagg	aaacatgttcc

*Synthesising fragment in Figure S4d*^1,2,3^

*Name	Template	Use	Cloning strategy	Sequence*

*surE* F	*A. baumannii*	Figure S4d construct	HiFi Assembly	tccacgctagatct*gaattc*ttttttcacaattttataatg

genome

*surE* R	*A. baumannii*	Figure S4d construct	HiFi Assembly	TCACAAAGTTTCTTTCTCGAcaaataaac

genome

*nlpD* F	*A. baumannii*	Figure S4d construct	HiFi Assembly	TCGAGAAAGAAACTTTGTGAataaag

genome

*nlpD* R	*A. baumannii*	Figure S4d construct	HiFi Assembly	ttgcgcactgacaa*aagctt*caaatcatttcggctaag

genome	agtaccaaaaaag**a**aagaggaaacatgttcc

Primers used to generate fragment containing the *araBAD* promoter, fused to *gfp*, with the *surE* terminator and *nlpD* promoter downstream, for cloning in pRW50 to make a *lacZ* *fusion in* *Figure 3e and 3f. Mutagenic primers can also be used with pSR*^1,2,3^

*Name	Template	Use	Cloning strategy	Sequence*

*araBAD* F	pBAD30	P*araBAD* section	HiFi DNA Assembly	ttgagtccacgctagatct*gaattc*atcgatgcataa

tgtgcctgtcaaatggac

*araBAD* R	pBAD30	P*araBAD* section	HiFi DNA Assembly	cctctttaatgctagcccaaaaaaacgggt

*araBAD* *nlpD*	genomic DNA	*nlpD* promoter section	HiFi DNA Assembly	ttgggctagcattaaagaggagaaattaag

GFP F

*araBAD* *nlpD*	genomic DNA	*nlpD* promoter section	HiFi DNA Assembly	ttgcgcactgacaa*aagctt*caaatcatttcggctaa

GFP R	gagtaccaaaaaag**a**aagaggaaacatgttcc

KO *surE* F	wild type	*surE* promoter mutation	Digestion-ligation	agatct*gaattc*ttttttcacaattttata**gc**gctatggag

gcataaatatat**gc**aattcaacaaataattc

KO *nlpD* R	wild type	*nlpD* promoter mutation Digestion-ligation	tgacaa*aagctt*caaatcatttcggctaagag**gc**cc

Aaaaaag**a**aagaggaaacatgttcc

*nlpD* improved	wild type	*nlpD* promoter mutation	Digestion-ligation	tgacaa*aagctt*caaatcatttcggctaa**tta**tacca

-10 element R	aaaaag**a**aagaggaaacatgttcc

^1^ Restriction endonuclease sites are italic and underlined

^2^ Mutations are in bold

^3^ *nlpD* reverse oligos have a single point mutation to inactivate a cryptic -10 element that becomes active upon mutation of the canonical *nlpD* promoter
